# Color constancy based on the geometry of color distribution

**DOI:** 10.1101/2020.05.19.105254

**Authors:** Takuma Morimoto, Takahiro Kusuyama, Kazuho Fukuda, Keiji Uchikawa

## Abstract

A white surface appears white under different lighting environments. This ability is referred to color constancy. The physical inputs to our visual system are dictated by the interplay between lights and surfaces, and thus for the surface color to be stably perceived, the illuminant influence needs to be discounted. To reveal our strategy to infer the illuminant color, we conducted three psychophysical experiments designed to test optimal color hypothesis: we internalize the physical color gamut under a particular illuminant and apply the prior to estimate the illuminant color. In each experiment, we presented 61 hexagons arranged without spatial gaps, where the surrounding 60 hexagons were set to have a specific shape in their color distribution. We asked participants to adjust the color of a center test field so that it appears a full-white surface placed under a test illuminant. Results and computational modeling suggested that although our proposed model is limited in accounting for estimation of illuminant intensity by human observers, it agrees fairly well with the estimates of illuminant chromaticity in most tested conditions. The accuracy of estimation generally outperformed other tested conventional color constancy models. These results support the hypothesis that our visual system can utilize the geometry of scene color distribution to achieve color constancy.

## 1. Introduction

The strength of cone excitations associated with a surface is determined both by the spectral composition of illuminant and by the surface spectral reflectance. Thus, when a scene illuminant changes, associated cone signals accordingly change. However, from the cone signals alone, we cannot work out how much of the change in cone signals is caused by the illuminant change, since the same change in cone signals could arise from a change in surface reflectance. Despite such difficulties, our perception of surface color of an object is remarkably stable under different lighting environments. For example, a white paper will appear white whether under bright sunlight, blue sky, or during a sunset. This visual function is often referred to as color constancy and enables us to identify objects by their color in spite of the change of illumination. However, the exact mechanisms underpinning this stability of the color vision system are not fully understood yet.

One common way to conceptualize color constancy is that the visual system first estimates the color of scene illumination based on cues available in the scene, and then discount its influence from the whole scene. Past color constancy research has been successful in identifying a number of mechanisms thought to underlie color constancy (Smithson, 2005; Hurlbert, 2007; Foster, 2011). In terms of neural substrates, both local (von Kries 1905; Smithson and Zaidi, 2004) and global (Ives, 1912; Werner, 2014) retinal adaptation is known to be useful in the implementation of color constancy. Identifying statistics-based cues to the illuminant has been a primary focus in behavioral color constancy studies. Perhaps the simplest method would be to find the brightest surface in the scene. This is based on the observation that a white surface with a completely flat spectral reflectance reflects any lights in a spectrally unmodified way, meaning that it conveys direct information about the color of the illumination (Land, 1977). Also, Tominaga, Ebisui, and Wandell (2008) suggested from a computational point of view that brighter surfaces are better cues than darker surfaces. Another simple but powerful transformation would be to compute mean chromaticity across all surfaces in a scene and assume that it is a good estimate of the illuminant color (Buchsbaum, 1980). This algorithm stands on the idea that the average color across all objects in a scene is typically grey (grey-world hypothesis), and thus the deviation from grey can be assumed to be due to the influence of the illumination. It is also known that spatial mean cone signals, rather than mean chromaticity, would be a good predictor of an illuminant color (Khang and Zaidi, 2004). The notion behind the use of such global statistics is that illumination typically lights a scene globally. It is known to be particularly powerful when the scene contains many different surface spectral reflectances. Foster and Nascimento (1994) found that the cone ratio between neighboring surfaces in natural scenes tends to be constant under the change of illuminants, allowing our visual system to access a signal invariant across the illuminant change. Unlike color constancy studies in the machine vision domain (Arjan, Theo and van de Weijer, 2011), it is important for studies on *human* color constancy, such as the present study, to experimentally verify that candidate cues are actually used by human observers. This is simply because finding something that is in theory useful does not guarantee that our visual system can or does use it. Moreover, it is good to remind ourselves that we may not rely on a single specific mechanism and rather use a combination of various cues. Such a strategy is useful in realistic situations as a specific cue may not be available in every scene (Kraft, and Brainard, 1999).

Using global scene statistics (e.g. mean chromaticity) to estimate the color of illumination has an attractive simplicity. However, whilst they reasonably account for experimental data, one of the longstanding questions in the field of human color vision was how our visual system distinguishes a white scene illuminated by a reddish illumination from a reddish scene illuminated by a whitish illumination when both cases produce the same reddish mean chromaticity (Brown, 1994). If our visual system relies purely on a chromaticity-based solution, the estimation of an illuminant color for these two situations should both be the same red. Golz and MacLeod (2002) provided a solution to this question. Firstly, they theoretically showed that red surfaces tend to have higher luminance when the illumination gets reddish compared to when the illumination is white. In contrast, when the illumination is white this chromatic-dependent luminance shift does not occur. Therefore, by looking at the correlation between luminance and chromaticity of given surfaces in a scene, it is possible to work out the color of the illumination lighting the scene. Secondly, importantly they have shown psychophysically that human observers are able to use this cue to solve such an ambiguity in a biased surface color set (e.g. dominantly reddish surfaces). Although some studies argued that the use of luminance-chromaticity correlation might be limited (Granzier et al., 2005; Golz, 2008), how luminances distribute over various chromaticities seems to play a key role in the implementation of color constancy.

Statistical models such as Bayesian estimation (Brainard and Freeman, 1997; Brainard et al., 2006) or low-level linear models to recover surface spectral reflectance (Maloney and Wandell, 1986) were reported to be good candidate models of human color constancy. Interestingly, these theoretical models make use of statistical regularities of surface spectral reflectance and illuminant spectra that occur in the natural environment, implying that the human visual system utilizes such constraints. This also makes sense from a mathematical point of view, as in general ill-posed problems can be solved when we put in a sufficient number of assumptions.

Natural scenes that we see in daily life appear to contain a wide variety of color, and thus it is tempting to think that any color can exist in a scene. However, this is a false intuition. In any scene, the possible range of chromaticity and luminance is limited by spectral composition of the illuminant, in a similar way that a digital monitor has a limited color gamut. This gamut for surface colors under a particular illuminant can be visualized by optimal colors or more precisely termed optimal surfaces (MacAdam, 1935a; MacAdam, 1935b). Optimal color is defined as a surface that has only 0% and 100 % reflectances and has at most two abrupt spectral transitions between them. We can think of two types of optimal colors: one boots up at wavelength λ_1_ and boots off at λ_2_ (band-pass type) and the other boots off and up at λ_1_ and at λ_2_ (band-stop type), where λ_1_ < λ_2_. To give an example how optimal colors distribute and how it changes depending on the color temperature of illuminants, we prepared 102,721 optimal colors by changing λ_1_ from 400 to 720 nm with 1 nm and λ_2_ from λ_1_ to 720 nm with 1 nm for the two types of optimal colors.

Figure 1 shows a distribution of 102,721 optimal colors (also known as MacAdam’s limit) and 49,667 objects in the real world drawn from the standard object color spectra database for color reproduction evaluation (SOCS, ISO/TR 16066:2003) under the illuminants of 3000K and 6500K. The left panel shows L/(L+M) in MacLeod-Boynton (MB) chromaticity diagram (MacLeod, and Boynton, 1979) versus luminance distribution and the right panel shows S/(L+M) versus luminance distribution. For the calculation of cone excitations, we used the Stockman and Sharpe cone fundamentals (Stockman and Sharpe, 2000). Optimal colors do not exist in a real world, but it is a useful mathematical tool to allow us to see the upper boundary of surface colors under a particular illuminant. If we look at this from the other side, under a particular illuminant any chromaticity has a unique corresponding optimal color, and the luminance of the optimal color gives the theoretical upper limit at the chromaticity.

**Figure 1:**
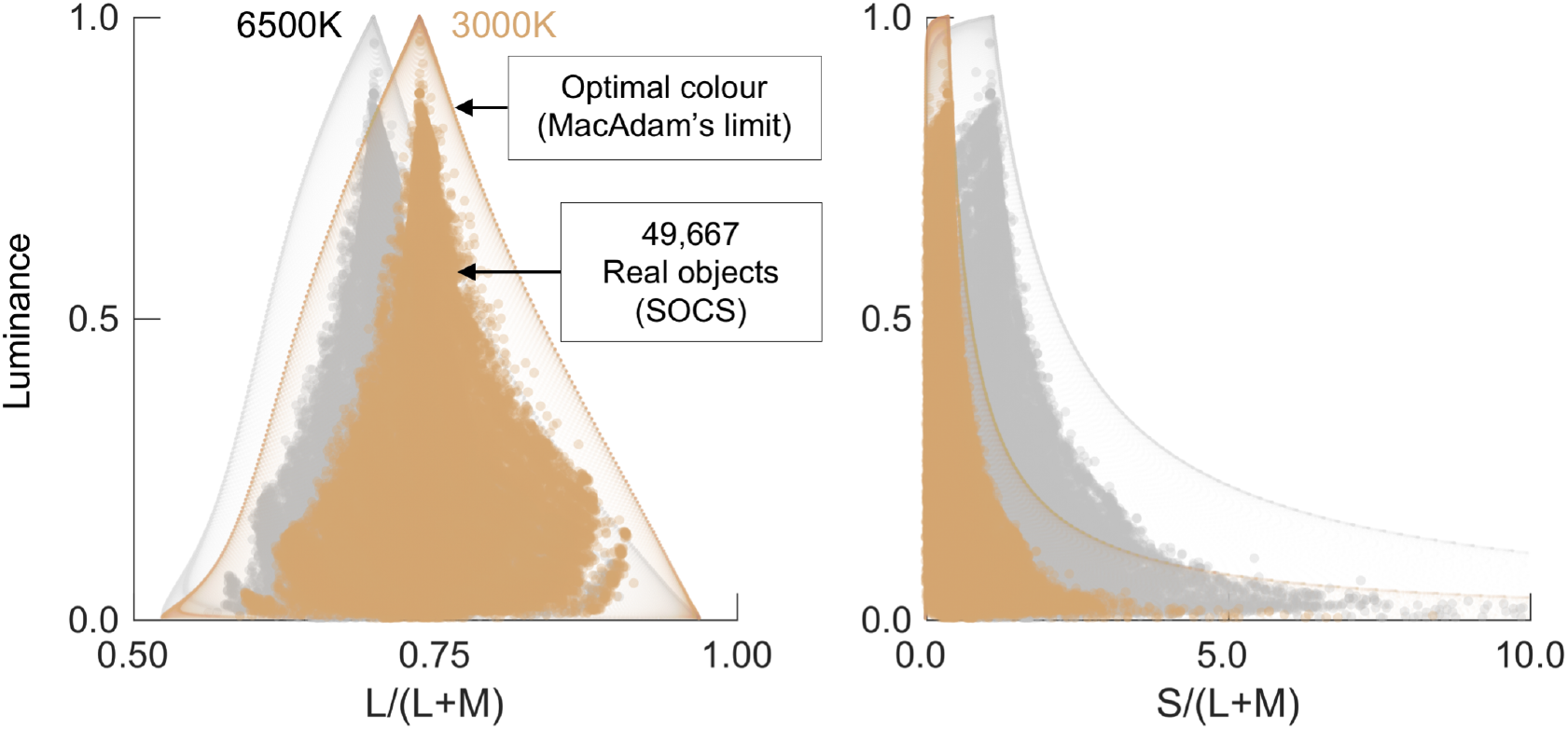
Color distribution of 102,721 optimal colors and 49,667 real objects in SOCS database. They are rendered under 6500K (gray distribution) and 3000K (orange distributions) to show the effect of illuminant change. The left and right panel show L/(L+M) vs. luminance and S/(L+M) vs. luminance distributions, respectively.

The peak of optimal color distributions always corresponds to a full-white surface (1.0 reflectance across all wavelengths), and thus it provides the chromaticity and intensity of the illuminant itself (i.e. white point). Optimal colors with higher purity have lower luminance, as they have a narrower-band reflectance the luminance distribution spreads out as the purity increases. We also see that the distribution of real objects is unsurprisingly fully included within the optimal color shell, but importantly their shape looks like the optimal color distribution under both illuminants. To put it in another way, there is a strong association between the illuminant color and the way a set of colors distribute in the real world.

This observation led to an idea: if our visual system is aware of this imposed statistical constraint, in other words if our visual system internalizes the shape of optimal color distribution under various illuminants, we can inversely work out from the observed chromaticity vs. luminance distribution to compute the influence of illuminant color. The simplest algorithm to implement this concept would be to find the best-fit optimal color shell to a given color distribution of a scene. This idea is depicted in Figure 2. To a given scene distribution, we fit the optimal color distribution of candidate illuminants (in this case (a) 3000K, (b) 6500K, (c) 20000K, (d) darker 20000K). For this example, the illuminant (c) fits the scene distribution best. The illuminants (a) and (b) are not appropriate as some surfaces exceed the optimal color distributions. Also, illuminant (d) does not hold some surfaces because the luminance level is too low even though the color temperature is the same as (c). This highlights the importance of selecting an appropriate illuminant intensity as well as a color temperature in the framework of optimal color model. Such an algorithm should function perfectly when a scene contains only optimal colors because we can uniquely find the optimal color distribution that fits to the scene distribution perfectly. However, in more general situations where a scene does not contain optimal colors, illuminant estimation is more challenging as we need to find a most appropriate optimal color distribution, for example by minimizing the root-mean-square-error between optimal color distribution and scene distribution.

**Figure 2:**
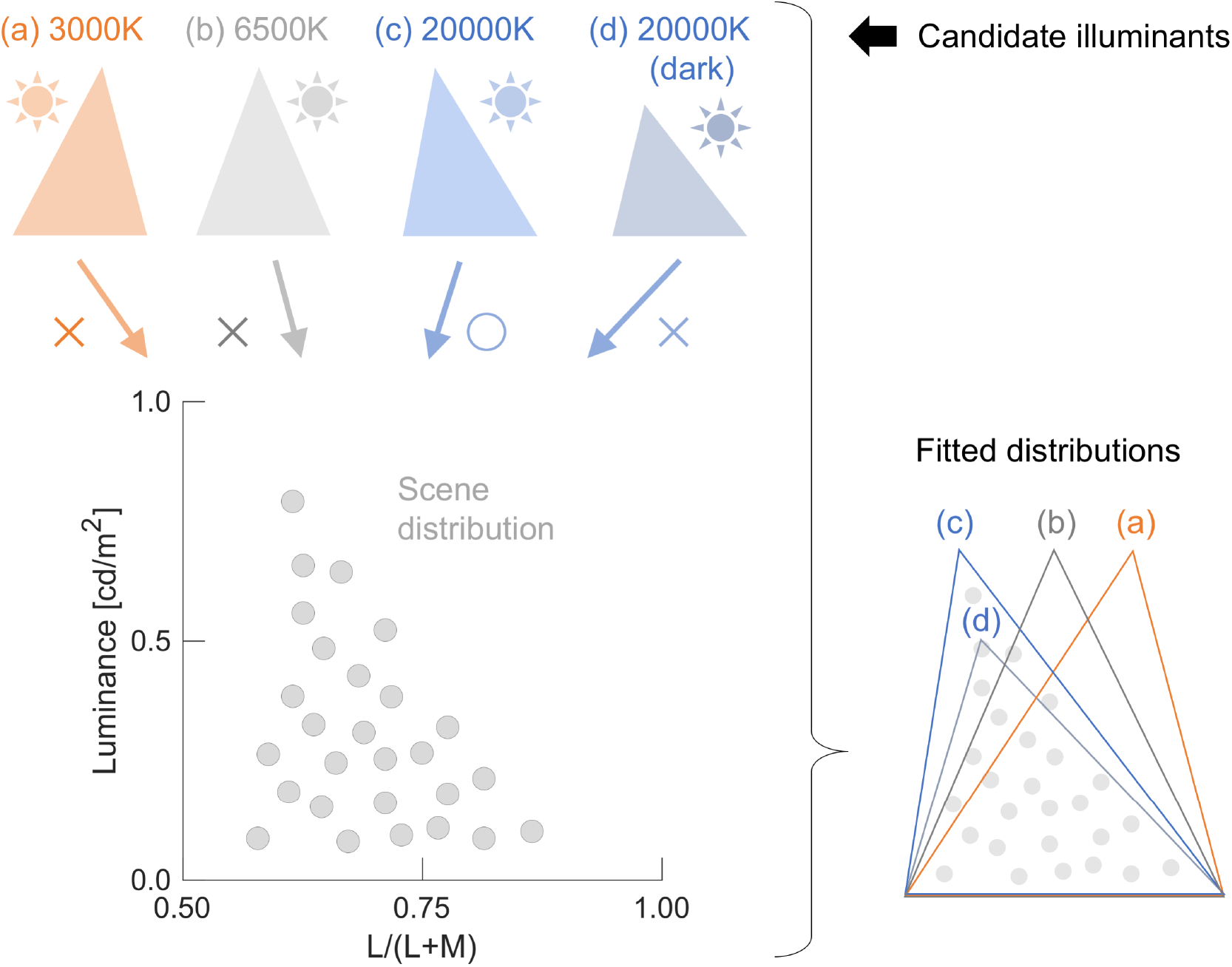
Schematic illustration of optimal color hypothesis: our visual system internalizes optimal color distributions and selects the one that gives a most likely fit to the scene distribution. In this case, (c) 20000K should be selected.

In summary, our hypothesis based on the optimal color model is as follows: when the visual system observes a scene, light sources and specular highlights have to be rejected first. Then, based on the remaining surfaces colors, our visual system seeks the most likely optimal color distribution such that it covers all surface colors in a scene. A computational algorithm based on an idea that utilizes the shape of RGB color distribution was also suggested (Forsyth, 1990).

In a series of previous papers, we have demonstrated the importance of luminance-chromaticity association (Uchikawa et al. (2012) and Morimoto, Fukuda, and Uchikawa (2016)). It was also shown that observers seem to ignore surfaces that appear self-luminous from the consideration of illuminant influence (Fukuda and Uchikawa (2014)). We also suggested that the optimal color model might describe bi-stable perception in the #TheDress phenomenon (Uchikawa, Morimoto and Matsumoto (2017)). The purpose of the present study is to test whether optimal color hypothesis quantitatively accounts for human observers’ behavior in a wider variety of conditions. Our approach to address this question is to prepare scenes that have various shapes of color distributions and to investigate whether observers’ estimations of illuminants follows the prediction by a computational model based on our hypothesis. We conducted three psychophysical experiments designed based on the following two scenarios.

Experiments 1 and 2 used optimal colors for scenes. Then, we manipulated the degree to which the scenes contained optimal colors. As briefly mentioned above, if a scene contains optimal colors, the optimal color model should function perfectly as we are able to uniquely find the illuminant under which the optimal color distribution perfectly matches the scene distribution. Therefore, even if the shape of the color distribution changes by darkening some optimal colors in the scene, it is expected that our estimate of illuminant color should not change.

In contrast, we used natural objects in Experiment 3. We implemented a computational optimal color model that incorporated our hypothesis, which predicts an illuminant color from a given color distribution. Guided by this model, we manipulated the shape of the natural color distribution so that the model prediction agreed or disagreed with set illuminant colors (i.e. ground-truth). Under this experimental set-up, we investigated the degree to which the model predicts observers’ estimation of the illuminant.

In each experiment, we used experimental stimuli that consisted of 61 hexagons spatially packed without a gap, where the surrounding 60 hexagons were designed to have a different shape of color distributions. The observer’s task was to adjust the chromaticity and luminance of the center test hexagon so that it appeared to be a full-white surface under a test illuminant. Results showed that observers’ estimations of illuminant intensity were not well predicted by our model, but other models relying on simple luminance statistics did not predict the observers’ behavior, either. However, estimation of illuminant chromaticity agreed well between human observers and our model prediction. In other words, human observers’ chromaticity settings changed or did not change in a manner predicted by the optimal color model. Although the applicability of our model is limited in some conditions, just as other models are, experimental results support the idea that our visual system can utilize the geometry of the color distribution to estimate the influence of the illuminant.

## 2. General method

### 2.1 Apparatus

All experiments were computer-controlled and conducted in a dark room. Experimental stimuli were presented on a cathode ray tube (CRT) monitor (Sony, GDM-520, 19 inches, 1600 × 1200 pixels) controlled by ViSaGe (Cambridge Research Systems), which allowed 14-bit intensity resolution for each phosphor. We performed gamma correction using a ColorCAL colorimeter (Cambridge Research Systems) and spectral calibration with a spectroradiometer (PR-650, Photo Research Inc.). Viewing distance was kept constant by a chin rest positioned 114 cm from the CRT monitor. Observers viewed stimuli binocularly.

### 2.2 Experimental stimuli

#### 2.2.1 Scene geometry

We used 61 hexagons as shown in Figure 3. Each hexagon was 2° diagonally, and the whole stimulus subtended 15.6° width × 14.0° height. The center hexagon was used as a test field, and its chromaticity and luminance were adjustable. The remaining 60 hexagons were surrounding stimuli and were designed to have a specific color distribution which is detailed in each experimental section.

**Figure 3:**
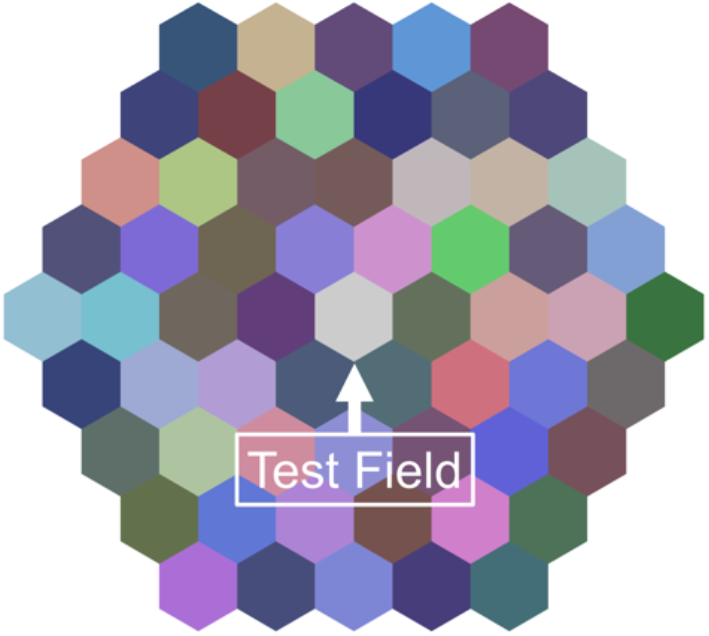
An example of an experimental stimulus configuration which consists of 61 uniformly-colored hexagons: the center test field and 60 surrounding stimuli (30 bright colors and 30 dark colors). Each hexagon subtended 2° diagonally. The chromaticity and luminance of the test field were adjusted by observers. This example is a *mountain* distribution under *6500K* in Experiment 1. The spatial pattern was shuffled for each trial.

#### 2.2.2 Test illuminants

We used 3000K, 6500K, and 20000K on the black-body locus as test illuminants for Experiments 1 and 2. For Experiment 3, we used 4000K, 6500K and 10000K.

### 2.3 Observers

Four observers (KF, KU, MS and TK; KF, KU and TK are the co-author of the study) participated in Experiment 1 and Experiment 2. Four different observers (HH, HY, RS and TM; TM is the first author of the study) were recruited for Experiment 3. Observers ages ranged between 22 and 63 (*mean* 31.5, *S.D.* 13.6). All observers had corrected visual acuity and normal color vision as assessed by Ishihara pseudo-isochromatic plates. RS and MS were naïve to the purpose of the study.

### 2.4 Implementation of optimal color model

Human observers infer the influence of illumination based on the shape of the chromaticity vs. luminance distribution in a scene. We implemented this idea based on a computational model that predicts the chromaticity and luminance of a scene illuminant from a set of 60 surface colors.

First, we normalized the luminances of the given 60 surfaces so that maximum luminance became 1.0. Thus, the 60 surfaces form a specific color distribution which peaks at 1.0, and we searched for an optimal color distribution that matches well the formed color distribution. We assumed that the model fully records the optimal color distribution (i.e. the chromaticities and the luminances of any possible optimal color) under 11,457 candidate illuminants: 57 color temperatures from 2000K to 30000K with 500 steps × 201 luminance levels from 1.0 to 3.0 (corresponding to the height of optimal color distribution) with 0.01 steps. Then, the model searches the optimal color distribution that is the best-fit to the observed color distribution, assessed by weighted root-mean-squared-error (*WRMSE*). If we take a given surface (*S_i_*), its luminance and the luminance of the optimal color at the chromaticity of *S_i_* can be written as *Lsi* and *Loi*, respectively. If we consider all surfaces from *S_1_* to *S_60_* in a scene, *WRMSE* for a specific candidate illuminant can be calculated by equation (1).

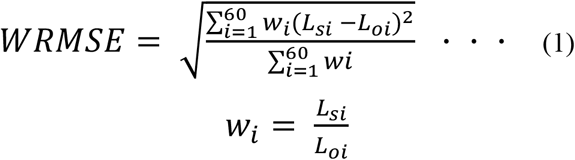

We weighted the luminance error by *w_i_* to put a greater weighting on brighter surfaces in proportion to the luminance of the corresponding optimal color. This is based on the finding that brighter colors have a greater influence on an observer’s estimation of the illuminant than darker colors (Uchikawa et al., 2012). Under some candidate illuminants, some surface colors might exceed the optimal color distribution, as in the case for illuminants (a), (b) and (d) in Figure 2. Thus, we first need to reject any illuminants under which any of 60 surfaces exceeds the optimal color shell (*Lsi* > *Loi* for any *i*). This restriction allows us to guarantee that we search only illuminants under which all surfaces are physically plausible (i.e. reflectance less than 1.0 at any wavelength). Then, we are looking for the illuminant from remaining candidates that minimizes the value of *WRMSE*. The chromaticity and the luminance of the best-fit illuminant gives the prediction of the optimal color model. Also note that we restricted our search along the black-body locus in this study since (i) we used test illuminants on the black-body locus and (ii) observers’ settings were also generally found on the locus.

### 2.5 Procedure

Before the first trial in each experiment, following one-minute dark adaptation, an observer first adapted for one minute to a white screen (2.85 cd/m^2^ equal energy white) covering the whole displayable area of the CRT monitor. Each experiment consisted of 9 conditions (3 test illuminants × 3 distributions), and thus 9 blocks formed one session. One block had five consecutive repetitions without inter-trial interval. The task of observers was to adjust the chromaticity and the luminance of the test field so that the test surface appeared as a full-white paper placed under a test illuminant. This is the so-called paper-match criterion introduced by Arend and Reeves (1986), and its nature is further argued by Reeves, Amano and Foster (2008). Additionally, our methodology here is a modified type of achromatic setting (Brainard, 1998), in which one response allows us to simultaneously measure the estimated illuminant chromaticity and intensity. Observers used a wired track ball (M570, Logitech) for two-dimensional adjustment of chromaticity in the MB chromaticity diagram and a number-pad for the adjustment of luminance. Both the initial chromaticity and luminance of the test field were selected randomly from a possible range for each trial. There was no time limitation. For each session, one distribution condition was randomly chosen first and fixed. Three illuminant conditions were tested for the distribution condition one after another in a random order. At the beginning of each block, observers adapted to the presented stimulus for 10 seconds (to be briefly adapted to a new illuminant) and then started the adjustment. They re-adapted to the full-white screen for 30 seconds between blocks. For each trial, the spatial arrangement of the 60 surrounding hexagons was shuffled. The observers in total performed two sessions in Experiment 1 and 2 and four sessions in Experiment 3. As a result, we collected 10 white point settings for each condition in Experiment 1 and 2 and 20 settings in Experiment 3.

## 3. Experiment 1

The purpose of the Experiment 1 was to investigate the effect of changing the shape of the chromaticity vs. luminance distribution on the estimated illuminant color. We used optimal colors for surrounding stimuli and manipulated the degree to which a scene contained optimal colors.

### 3.1 Color distribution of surrounding stimuli

In Experiment 1, we chose three sets of 60 surfaces (30 bright colors and 30 dark colors) that produce different chromaticity-luminance distributions: *mountain*, *reverse*, and *flat*. Dark colors always had the same chromaticity and 20% of the luminance of the bright colors. Figure 4 helps us to see how 30 chromaticities were chosen and how luminance values were assigned to each chromaticity. We first picked the chromaticities of three colors (labelled as Red, Green and Blue in Figure 4 (a)). The criteria to set these colors were that (i) they do not exceed the chromatic gamut of the CRT monitor under any test illuminant (*3000K*, *6500K*, and *20000K*) and that (ii) they have as high purity as possible. Then, we connected three colors by straight lines, which forms the magenta triangle.

**Figure 4:**
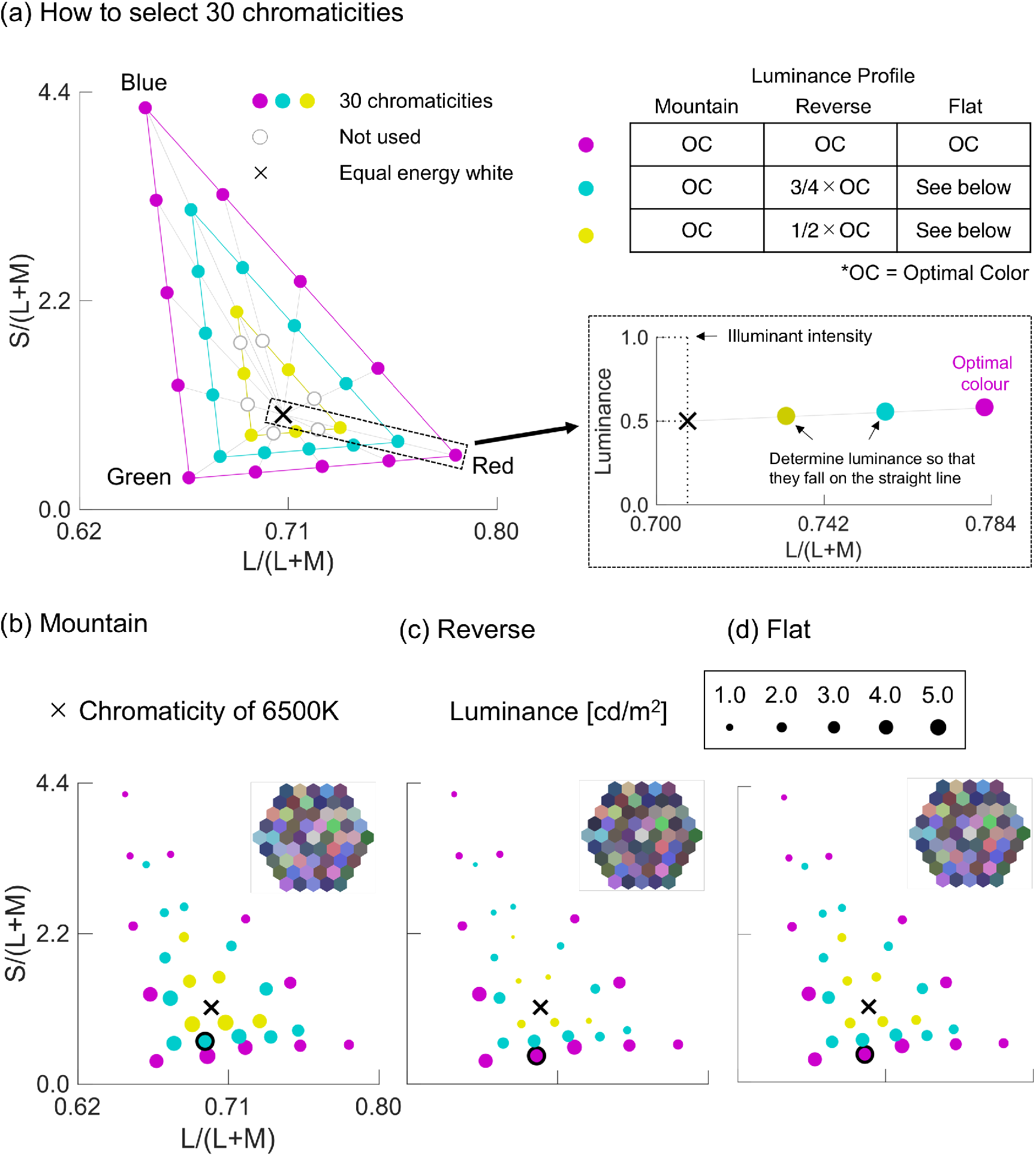
(a) How we selected 30 chromaticities and their luminance values for each distribution condition in Experiment 1. See main text for details. (b) - (d) Color distributions for each distribution condition. The symbol size expresses the luminance value at each chromaticity. The highest luminance color is marked by black edge. The black cross indicates the chromaticity of 6500K illuminant. Note only the *6500K* condition is shown. Also, only the 30 bright colors are presented here.

We picked three colors on each side so that the gaps between colors have equal distance. Next, we drew cyan and yellow triangles so that they have similar figures to that of the magenta triangle (2:3 and 1:3, respectively) and again picked colors on each side in the same way. As shown by white circles on the yellow triangle, we excluded some colors simply to adjust the number of chromaticities to 30. As a result of this selection, we have three purity levels shown by magenta, cyan and yellow triangles, and each level had 12, 12, and 6 colors, respectively. Here we note that higher purity colors are more informative regarding the illuminant color according to the framework of optimal color hypothesis. For example, when a scene has a saturated bright reddish surface, the illuminant is unlikely to be bluish as it cannot cover the surface.

Next, we assigned luminance values for each of 30 chromaticities by defining surface spectral reflectance for each chromaticity. An inserted table at the top-right corner in Figure 4 (a) shows how luminances were chosen for each distribution condition (*mountain*, *reverse* and *flat*). Figure 4 (b), (c) and (d) show resultant distributions for each distribution condition under *6500K*, where the symbol size indicates the luminance for each chromaticity.

For the *mountain* distribution all colors were set to optimal colors. Note that the optimal color distribution peaks at an illuminant’s chromaticity, and the luminance decreases as it gets away from the white point. Thus, for the *mountain* condition, luminance values were always highest at yellow symbols and decreased towards magenta symbols as shown in Figure 4 (b). We also note that it is generally not possible to uniquely convert a chromaticity to a surface spectral reflectance simply because there are myriad of potential reflectances that could produce the desired chromaticity. However, here we only considered optimal surfaces, allowing us to uniquely find a surface spectral reflectance that produces a desired chromaticity under equal energy white. In this way, 30 chromaticities were converted to spectral reflectances of optimal colors, with which we simulated the effect of an illuminant change to *3000K*, *6500K* and *20000K*.

*Reverse* and *flat* conditions were prepared by manipulating the spectral reflectance functions of the *mountain* distribution. For both the *reverse* and *flat* distribution, stimuli with chromaticities on the magenta triangle were again set to optimal surfaces. However, for the *reverse* distribution, stimuli with chromaticities on the cyan triangle were set to have three-fourths of the reflectance of optimal surfaces at the corresponding cyan chromaticities (i.e. 0.75 and 0.00 reflectance across wavelengths), and stimuli with chromaticities on yellow triangle were set to have half of the reflectance of optimal surfaces at the corresponding yellow chromaticities (i.e. 0.50 and 0.00 reflectance across wavelengths). As a result, unlike the *mountain* distribution, luminance values in the *reverse* condition were highest at the magenta symbols and decreased towards the white point (black cross) as shown in Figure 4 (c). For the *flat* distribution, the bottom-right subpanel in Figure 4 (a) helps us to understand our manipulation. The luminance values at the cyan and yellow chromaticities were determined so that they fall on the straight line connecting the luminance of the optimal color of the magenta triangle at half of the illuminant intensity. As shown in Figure 4 (d), the resultant distribution appears somewhere between the *mountain* and *reverse* conditions, showing relatively flat luminance values over all chromaticities.

We emphasize that chromaticity values for each surface did not change at all depending on the distribution condition, and thus any chromaticity-based illuminant estimation algorithm, such as the mean chromaticity model, predicts exactly the same illuminant color for each distribution. This was held for all illuminant conditions.

After the selection of surface reflectances, we adjusted the intensity of test illuminants so that mean luminance across 60 colors becomes 1.2 cd/m^2^ for all conditions. As a result, illuminant intensities were set to 2.96 cd/m^2^, 2.97 cd/m^2^, and 2.97 cd/m^2^ for *mountain*, *reverse*, and *flat* distributions in the *3000K* condition, respectively. For *6500K*, they were 4.14 cd/m^2^, 4.16 cd/m^2^, and 4.14 cd/m^2^. Finally, for *20000K*, we used 3.48 cd/m^2^, 3.49 cd/m^2^, and 3.48 cd/m^2^.

### 3.2 Results and discussion

Panel (a) in Figure 5 shows the averaged observers’ settings across 10 trials in a MB chromaticity diagram. Error bars are not shown to increase the visibility. The shape of symbol indicates the distribution condition (*mountain*, *reverse*, and *flat*), and colors indicate the illuminant conditions (*3000K*, *6500K*, and *20000K*). Different sub-panels show the results for different observers. Each colored cross symbol indicates the chromaticity of the test illuminant, and thus they show the ground-truth in each illuminant condition.

**Figure 5:**
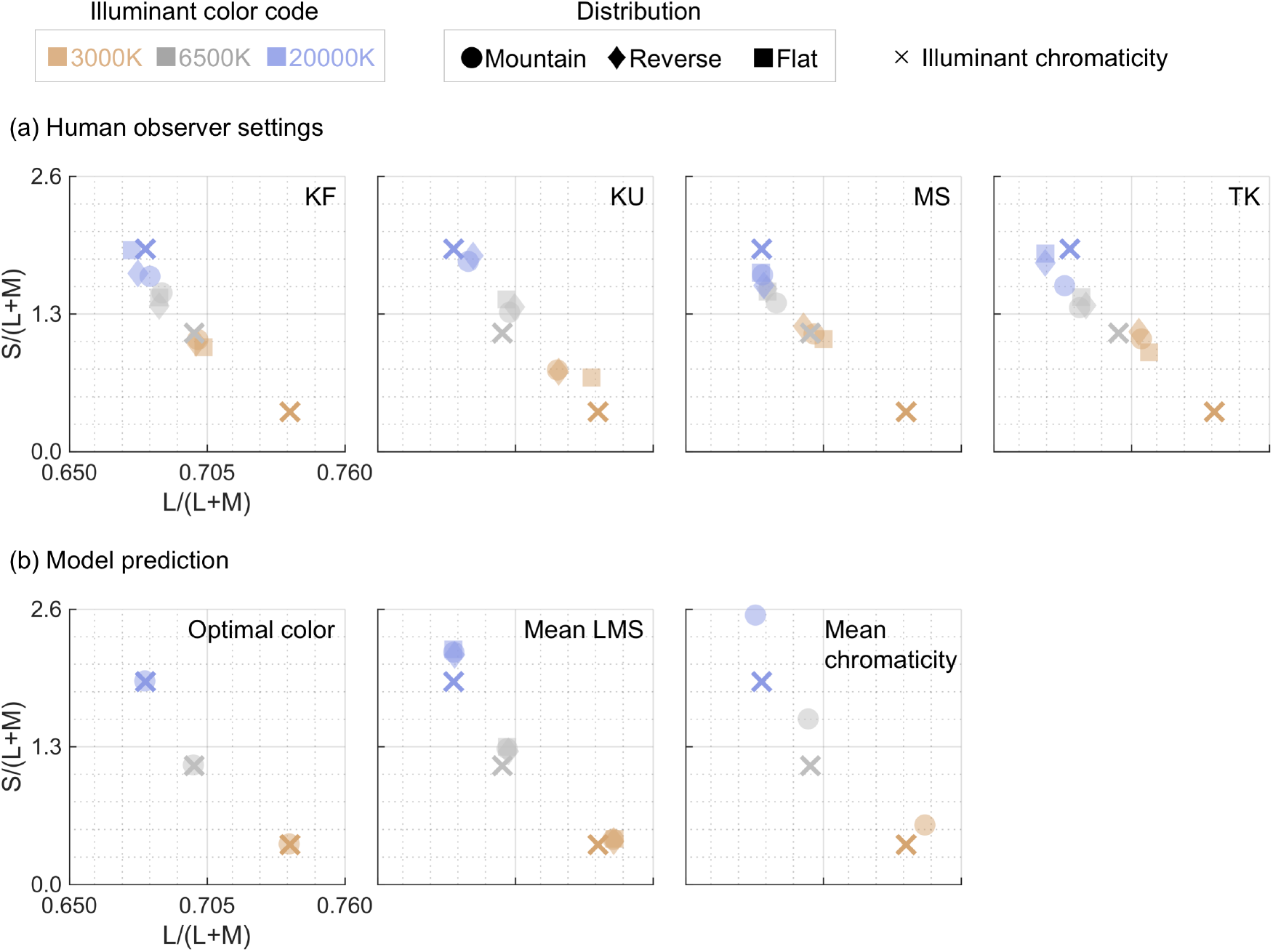
(a) Observer’s settings plotted on a MB chromaticity diagram for each condition in Experiment 1. The shape of each symbol indicates distribution condition, and color indicates illuminant condition. The colored cross symbols show chromaticities of the test illuminants. Different sub-panels correspond to different observers. (b) Model prediction by optimal color model, mean LMS model and mean chromaticity model. Note that the optimal color model and mean chromaticity predict the same chromaticity for any distribution condition by experimental design, and thus only the prediction for the *mountain* condition is shown for the sake of clarity.

Panel (b) shows the predictions from the optimal color model, mean LMS model and mean chromaticity model. Note that in this experiment the optimal color model provides a perfect estimation of illuminant chromaticity regardless of the distribution condition due to the experimental design. For this reason, we plotted only predictions for the *mountain* condition. This is self-evident as we used optimal colors for stimuli in this experiment, but we also applied fitting procedures to each of 9 conditions based on equation (1) for a sanity check. This confirmed that predicted chromaticity and intensity for each condition indeed matched the chromaticity and the intensity of a test illuminant. To calculate the prediction of the mean LMS model, we first averaged cone responses across the 60 surrounding hexagons, and then we converted the averaged cone responses to MB chromaticity coordinates. Note that this manipulation is equivalent to calculating the luminance-weighted average of MB chromaticity coordinates. Here, it is shown that estimated chromaticities by mean LMS are positioned very close to the chromaticity of test illuminants, and also there is little difference across distribution conditions under any illuminant condition. The mean chromaticity model provides prediction based on the average chromaticity across the 60 surrounding hexagons. Their predictions are deviated mainly toward higher S/(L+M) direction from the illuminant chromaticities, and it does not change depending on the distribution condition. This is why only predictions for *mountain* condition is provided in the figure.

If an observer is perfectly color constant, observers’ settings in panel (a) should superimpose on the chromaticity of the corresponding test illuminant shown by cross symbols. However, that was never the case in this experiment. Instead all settings showed some deviation from the ground-truth points, and some systematic trends are observed as follows.

First, for all observers we see that observers’ settings for the three distribution conditions are rather closely clustered. This trend also roughly holds for any illuminant condition. These suggest that the shape of chromaticity-luminance distribution has little impact on observers’ estimation of the illuminant chromaticity in this experiment, which is predicted by the optimal color model, mean LMS model and mean chromaticity model as shown in panel (b).

Second, as predicted, observers’ settings are separated depending on the illuminant condition. Thus, unsurprisingly, the color temperature of the test illuminant has a strong effect on observers’ settings. The degree of separation across the illuminant condition appears to depend on observers. For example, settings for MS are closely gathered across test illuminants while KU shows a fairly greater separation.

Our methodology allowed observers to simultaneously adjust luminance in addition to chromaticity, which gives us the indication of an observer’s estimation of illuminant intensity. Figure 6 shows the luminance setting for each condition, averaged across four observers. To see whether observers’ settings show agreement with simple luminance statistics, red and cyan crosses indicate the mean luminance and the highest luminance across the 60 surrounding surfaces, respectively. Blue crosses show the intensities of test illuminants (i.e. ground-truth), and thus if an observer’s setting matches blue crosses, it indicates that the observer perfectly estimated the intensity of the illuminant. Note that in this experiment the optimal color model provided the perfect estimation of illuminant intensity for all conditions by design, and thus the blue cross symbols also indicate the prediction of the optimal color model.

**Figure 6:**
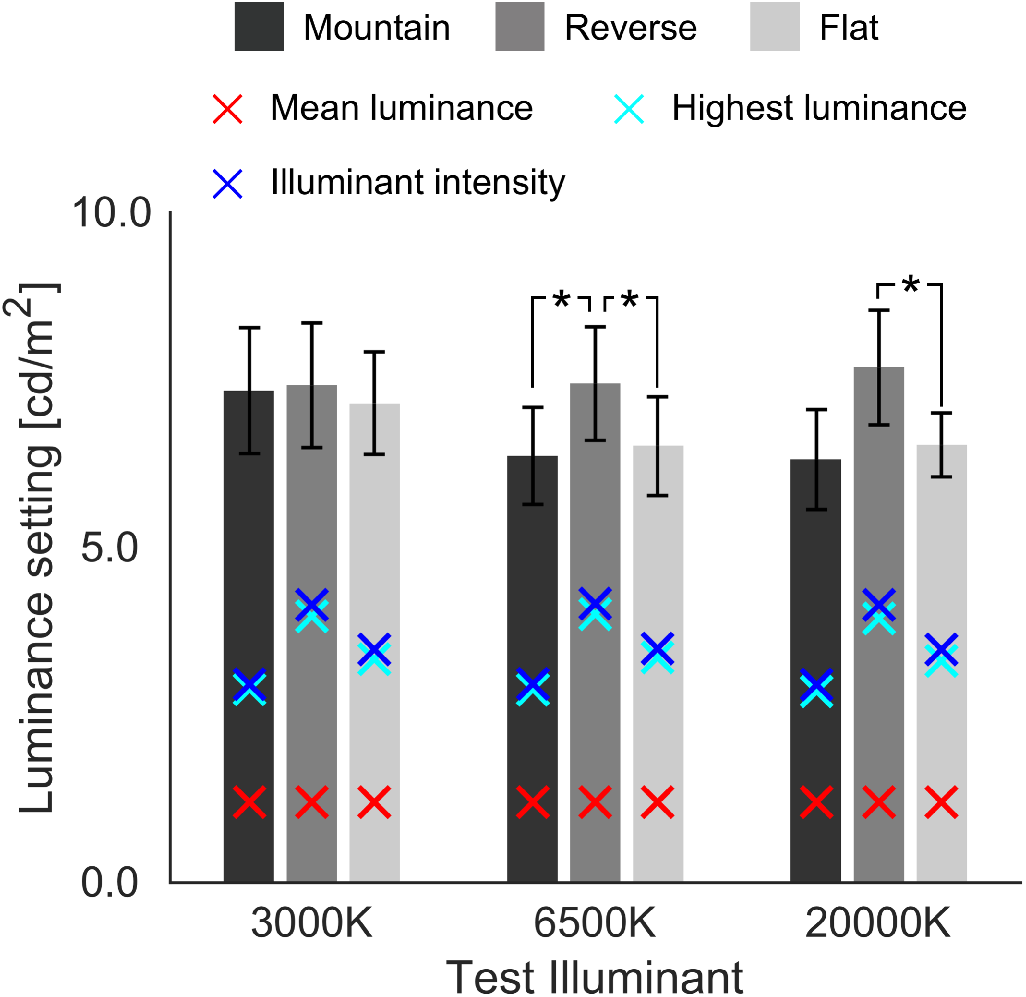
Observer’s luminance settings for each condition in Experiment 1. Distribution condition is labelled by the lightness of the bars. Red cross symbols indicate the mean luminance across the 60 surrounding surfaces, and the cyan symbol is the highest luminance of the 60 surfaces. Blue cross symbols show the intensity of the set test illuminant and therefore indicate the prediction by the optimal color model. The error bars indicate ± S.E. across four observers. Asterisks (*) show a significant difference (α < 0.05, Bonferroni’s correction).

However, observers’ estimations of intensity were never perfect. Rather, their settings generally exceeded the set illuminant intensity, showing observers assumed the illuminants to be much more intense than they actually are.

To quantitively evaluate whether the pattern of luminance settings can be explained by simple luminance statistics, we calculated correlation coefficient between observers’ average settings and each luminance statistic across 9 conditions. Correlation coefficients were 0.0788 (*p* > 0.05), 0.710 (*p* = 0.0320) and 0.700 (*p* = 0.0359) for mean luminance, highest luminance, and illuminant intensity, respectively. Thus, the highest luminance model and illuminant intensity (i.e. prediction from the optimal color model) showed a significant correlation. These do not rule out that observers used different statistics or a more complicated strategy, but it suggests that their behaviors can be explained reasonably well by the optimal color model or simple luminance statistics (highest luminance in this case).

Next, a two-way repeated-measures analysis of variance (ANOVA) was performed for **(a) distribution condition** (*mountain*, *reverse* and *flat*) and **(b) illuminant condition** (*3000K*, *6500K* and *20000K*) using within-subject factors for the luminance settings. The main effect of **(a) distribution condition** and **(b) illuminant condition** were not significant (*F*(2,6) = 3.48, *p* > 0.05; *F*(2,6) = 3.88, *p* > 0.05). However, the interaction between the two factors was significant (*F*(4,12) = 5.45, *p* = 0.00975).

Further analysis of the interaction revealed that the simple main effect of **(a) distribution condition** was significant at *6500K* and *20000K* (*F*(2,6) = 9.71, *p* = 0.0132; *F*(2,6) = 7.80, *p* = 0.214, respectively), but not at *3000K* (*F*(2,6) = 0.15, *p* > 0.05). Furthermore, the simple main effect of **(b) illuminant condition** was significant for *mountain* condition (*F*(2,6) = 5.48, *p* = .0443), but not significant for *reverse* and *flat* (*F*(2,6) = 1.39, *p* > 0.05; *F*(2,6) = 3.76, *p* > 0.05, respectively).

To further clarify the relation across levels regarding the simple main effect of **(a) distribution condition** and **(b) illuminant condition**, the results of multiple comparisons with Bonferroni’s correction (significance level *α* = 0.05) where significant differences were found are shown by the asterisk (*) in the Figure 6. Overall, the results of these statistical analysis suggest that luminance settings were (i) higher at *reverse* than *mountain* and *flat* for *6500K* and (ii) higher at *reverse* than *flat* for *20000K*.

Next, to argue how much color constancy held for each condition, it would be helpful to quantify the degree of constancy. Several metrices were used in past studies (summarized in Foster (2011)), but a fundamental idea is to capture how much observers’ subjective white p€oints shift in relation to the shift of physical change of illuminant chromaticity. In this study, we used an index that considers both the direction and the amount of the observers’ settings shift.

As shown in Figure 7 we first defined two vectors 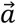 and 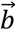, where 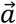 is a vector that originates from the chromaticity of illuminant *6500K* to the illuminant chromaticity of *3000K* or *20000K*. The vector 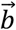 extends from the observers’ settings under 6500K to settings under the test illuminant (*3000K* or *20000K*). We defined *θ* as indicating the angle that two vectors create, as shown in right inserted panel in Figure 7. Constancy index (CI) is calculated by equation (2).

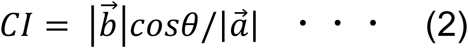

**Figure 7:**
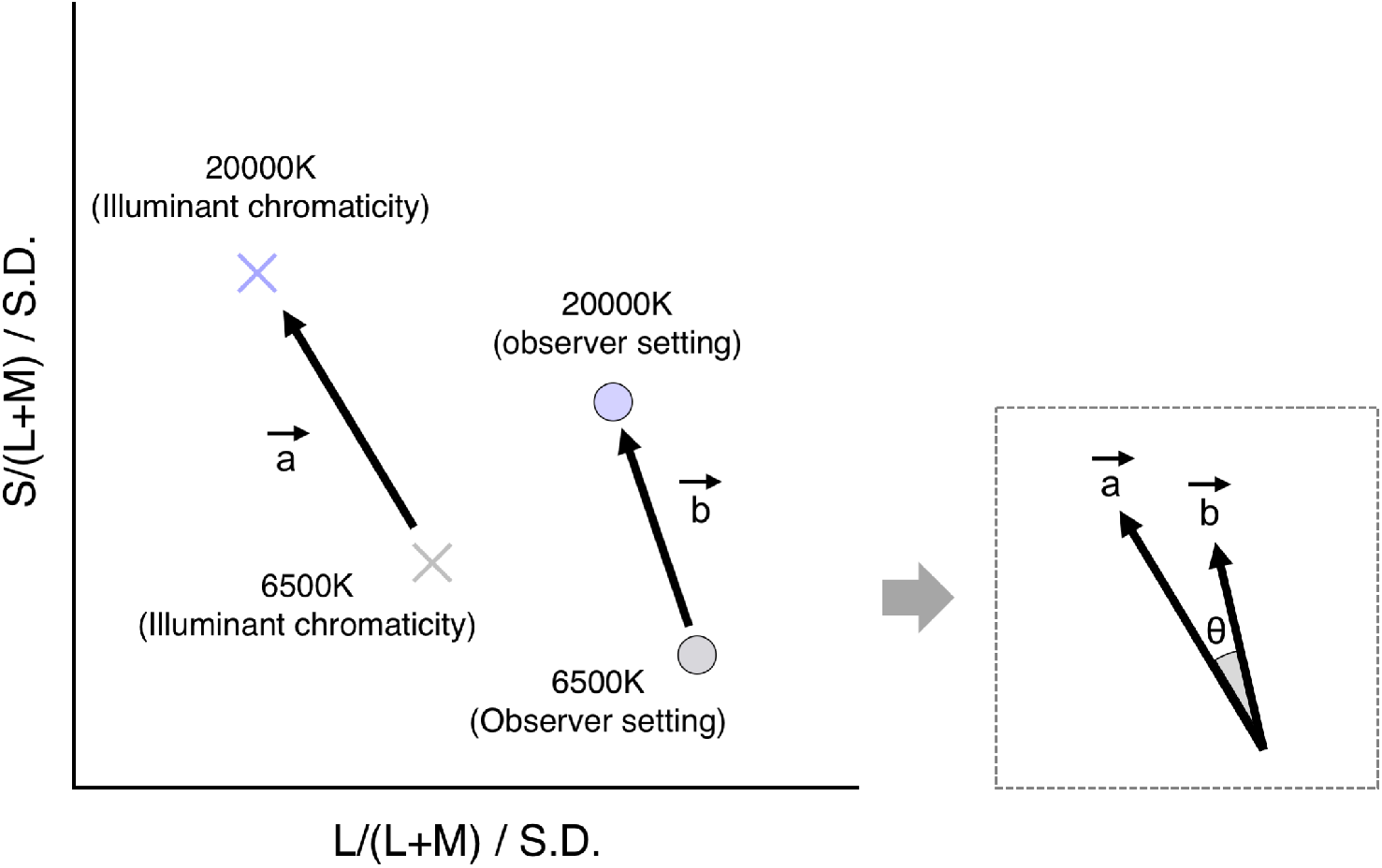
How to define constancy index (*CI*). Vectors 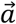 and 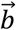 show the shift of physical illuminant chromaticity and the shift of perceptual white point, respectively, and *θ* is the angle between the two vectors. Note that each axis was divided by the average of standard deviations across 4 observers for the *mountain*-*6500K* condition to compensate for the scale difference along L/(L+M) and S/(L+M). *CI* is defined by 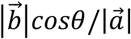 in this scaled MB chromaticity diagram.

If a shift of an observer’s setting caused by illuminant change is the same as a shift of illuminant chromaticity in terms of both distance and direction, *CI* becomes 1.0. Any deviation in distance or direction would lower the *CI*. In theory *CI* can take a value more than 1.0 when 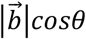 is larger than 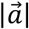, but we did not find such a case in this study. If the direction of shift perfectly matches between 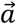 and 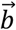 (i.e.*θ* = 0), cos*θ* becomes 1.0 and thus it does not lower the *CI* value. In this sense, cos*θ* serves as a loss factor due to the directional deviation.

Also note that a problem with calculating a distance-based metric in MB chromaticity diagram is that scales for L/(L+M) and S/(L+M) are different and arbitrary. Thus, to give approximately equal consideration to both directions, we divided each axis by the average of standard deviations across four observers of the settings under the *mountain*-*6500K* condition (which we consider as the standard condition). All *CIs* were calculated in this scaled MB chromaticity diagram.

The bar chart in Figure 8 shows averaged *CI* across 4 observers for each condition. First it was shown that *CIs* were roughly around 0.5 for the *mountain* and *reverse* conditions for *3000K* and *CIs* were slightly lower for *20000K*. The *flat* condition has a slightly higher *CI*, especially for *3000K*.

**Figure 8:**
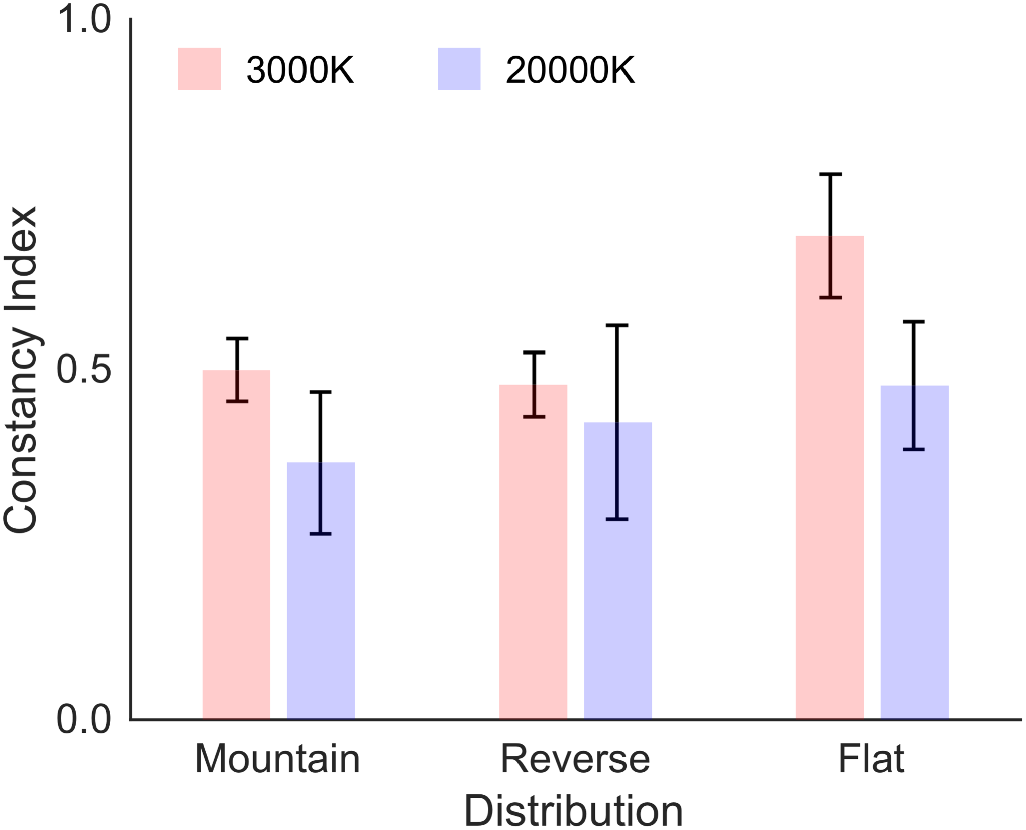
Constancy index (*CI*) averaged across four observers in Experiment 1. Note that *CIs* closer to 1.0 indicates better color constancy. Error bars indicate ± *S.E.* across 4 observers.

We performed a two-way repeated-measures ANOVA for **(a) distribution condition** (*mountain*, *reverse* and *flat*), and **(b) illuminant condition** (*3000K* and *20000K*) as within-subject factors for the *CIs*. The main effect of **(a) distribution condition** was significant (*F*(2,6) = 8.40, *p* = 0.0182) while the main effect of **(b) illuminant condition** and the interaction between two factors were not significant (*F*(1,3) = 7.63, *p* > 0.05; *F*(2,6) = 0.59, *p* > 0.05, respectively).

Multiple comparisons with Bonferroni’s correction (significance level *α* = 0.05) for **(a) distribution condition** found that the *flat* condition showed higher *CIs* than the *mountain* and *reverse* conditions, but there was no significant difference between the *mountain* and *reverse* conditions.

Overall, the statistical analysis above indicated that color constancy works equally well under *3000K* and *20000K*, and also it works better for the *flat* distribution than other distributions in Experiment 1.

We have so far argued the degree to which human observers achieved color constancy. However, the main purpose of the present study is to quantify how well our optimal color models accounts for the pattern of human observers’ settings, ideally in comparison with other candidate color constancy models. The *CI* calculates how much an observer’s setting shifted in relation to the shift of the physical illuminant chromaticity. Thus it gets higher when the shifts of perceptual white point and physical illuminant are close to each other in distance and direction, and it becomes 1.0 when the two shifts perfectly match. Although *CI* was originally designed to reflect the degree of color constancy, it can also be considered as a metric which indicates how much physical shift of illuminant chromaticities predict observers’ settings. Based on this idea, we quantified the degree to which our optimal color model and other computational models (mean LMS and mean chromaticity) predicted human observers’ settings. We introduce model index (*MI*) as defined by equation (3), which replaces the x symbols in Figure 7 to the + symbols indicating predictions from a model as shown in Figure 9.

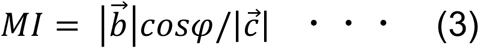

**Figure 9:**
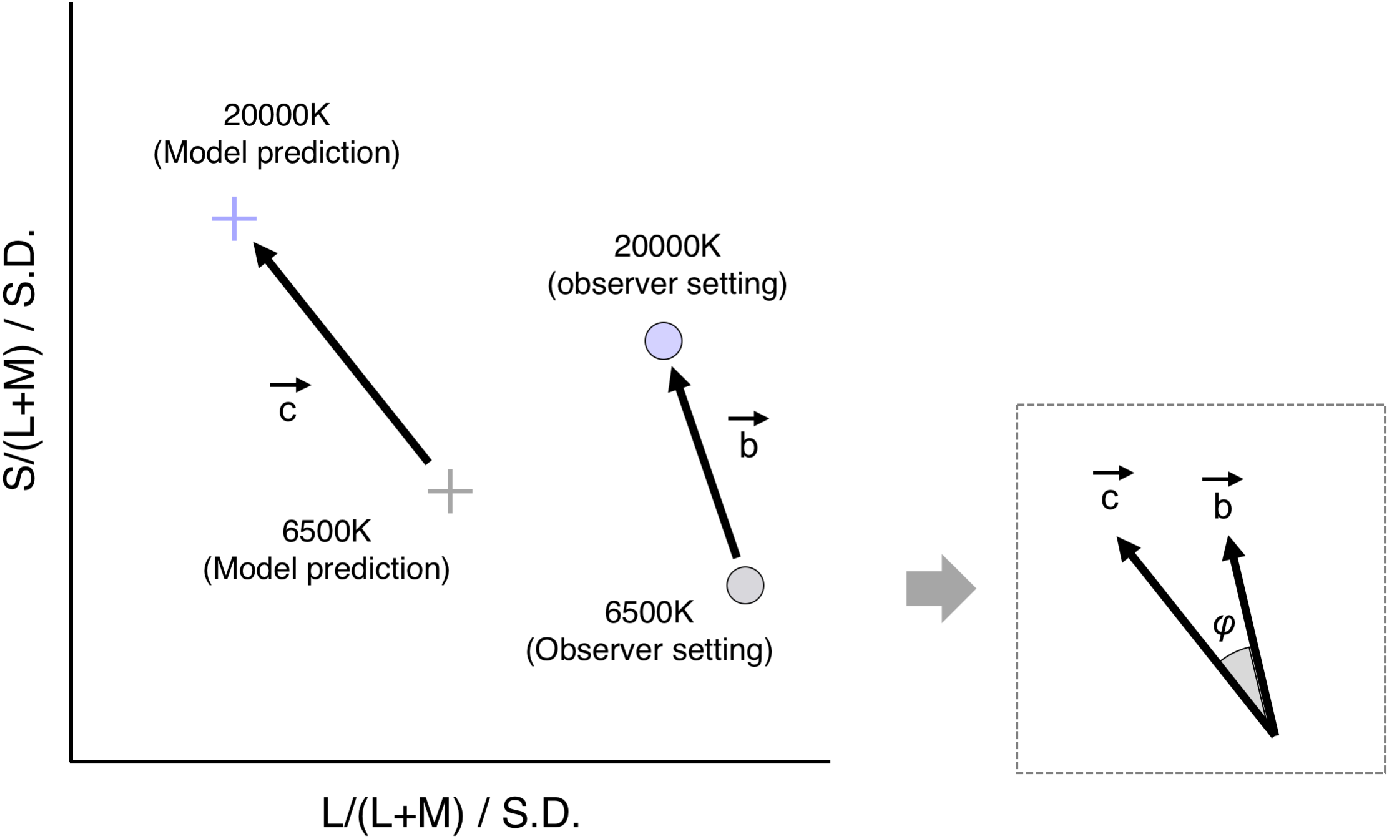
How to define model index (*MI*) which quantifies the degree to which three models of interest (optimal color, mean LMS, and mean chromaticity) account for human observers’ settings. Vectors 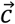 and 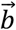 show the shift of chromaticities predicted by a model and the shift of perceptual white point, respectively. *φ* is the angle between two vectors. *MI* is defined by 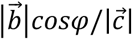 in the scaled MB chromaticity diagram.

Higher values indicate better model prediction and 1.0 means perfect prediction. Figure 10 shows *MIs* in each condition. Note that in this experiment the optimal color model predicted illuminant chromaticity perfectly and therefore the values of *MIs* match those of *CIs*. The mean LMS model first averaged cone signals across 60 surfaces and the average cone signal was transformed to MB chromaticity coordinates. The mean chromaticity estimates illuminant color based on the average chromaticity across 60 surfaces.

**Figure 10:**
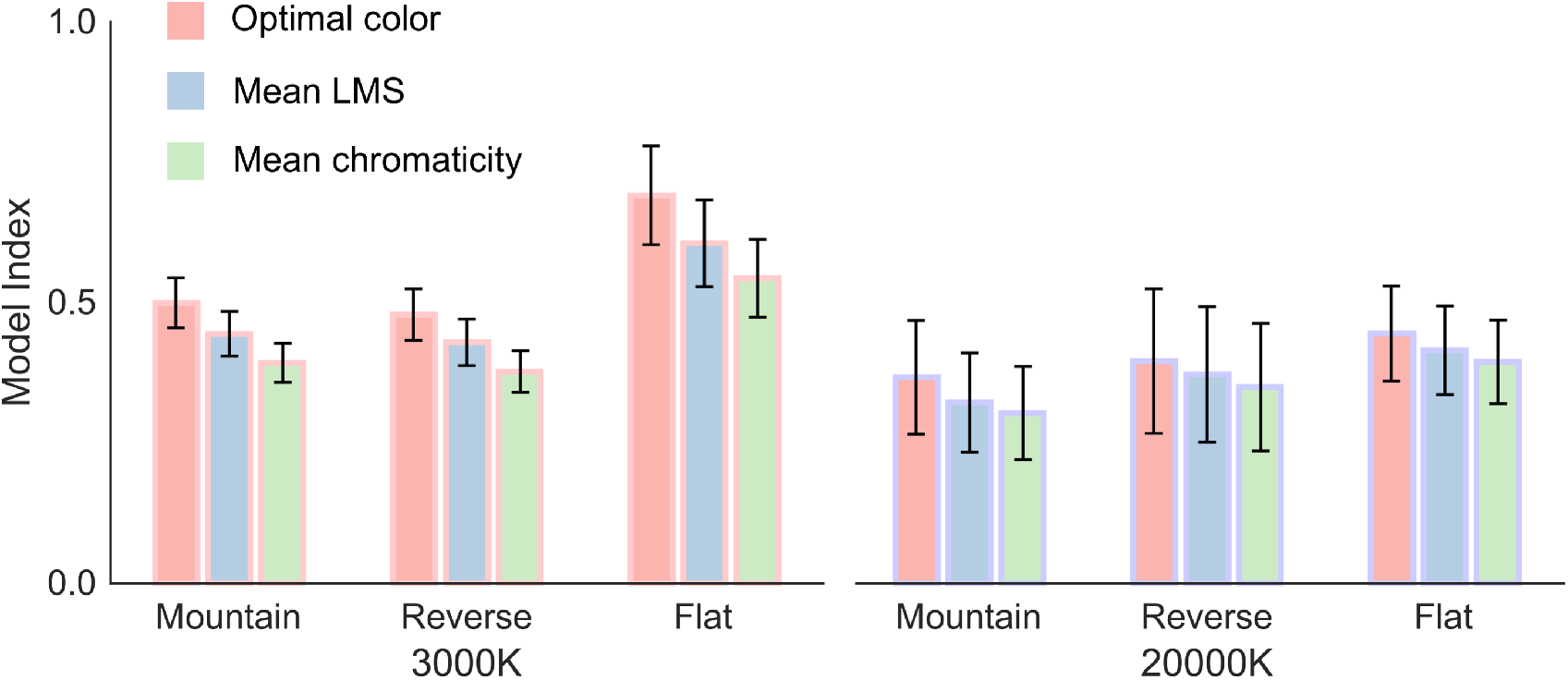
Model index (*MI*) for each condition in Experiment 1. *MIs* were calculated separately for each observer first and then averaged across 4 observers. Higher *MI* indicate that a model predicted human observers’ settings better. Error bars show ±*S.E.* across 4 observers.

We see that *MIs* values are high in the order of optimal color, mean LMS, and mean chromaticity models for all conditions. We ran a following analysis to confirm if our model prediction was statistically better than mean LMS and mean chromaticity models.

We performed a three-way repeated-measures ANOVA for **(a) model type** (*optimal color*, *mean LMS* and *mean chromaticity*), **(b) distribution condition** (*mountain*, *reverse* and *flat*), and **(c) illuminant condition** (*3000K* and *20000K*) as within-subject factors for the *MIs*. The main effect of **(a) model type** and **(b) distribution condition** were significant (*F*(2,6) = 52.8, *p* = 0.000155; *F*(2,6) = 7.93, *p* = 0.0207, respectively) while the main effect of **(c) illuminant condition** was not significant (*F*(1,3) = 8.51, *p* > 0.05). The interaction between **(a) model type** and (**b) distribution condition** and between **(a) model type** and **(c) illuminant condition** were significant (*F*(4,12) = 15.4, *p* = 0.000113; *F*(2,6) = 220.9, *p* < 0.00001, respectively). In contrast, the interaction between **(b) distribution condition** and **(c) illuminant condition** was not significant (*F*(2,6) = 0.57, *p* > 0.05). The interaction among three factors was not significant, either (*F*(4,12) = 3.11, *p* > 0.05).

We analyzed simple main effects for the interaction between **(a) model type** and **(b) distribution condition**, and following results were shown. The simple main effect of **(a) model type** was significant for all distribution conditions (*F*(2,18) = 45.4, *p* < 0.00001 for *mountain*; *F*(2,18) = 42.9, *p* < 0.00001 for *reverse*; *F*(2,18) = 62.8, *p* < 0.00001 for *flat*). Also the simple main effect of **(b) distribution condition** was significant for all models (*F*(2,18) = 8.36, *p* = 0.00271; *F*(2,18) = 7.74, *p* = 0.00375; *F*(2,18) = 7.69, *p* = 0.00386, respectively). A post-hoc multiple comparison using a Bonferroni’s correction (significance level, 0.05) revealed a following relation: (i) *mountain* = *reverse*, *mountain* < *flat*, and *reverse* < *flat* for *optimal color model*, and (ii) *mountain* = *reverse*, *mountain* < *flat*, and *reverse* = *flat* for *mean LMS* and *mean chromaticity models*. We also found following results: *optimal color model* > *mean LMS model*, *optimal color model* > *mean chromaticity model*, and *mean LMS model* > *mean chromaticity model* for all distribution conditions (i.e. *mountain*, *reverse* and *flat* distributions).

Next, we analyzed simple main effects of interaction between **(a) model type** and **(c) illuminant condition**. We found that **(a) model type** showed significant simple main effects both at *3000K* and *20000K* (*F*(2,12) = 97.7, *p* < 0.00001; *F*(2,12) = 19.8, *p* = 0.000158, respectively). Moreover, the simple main effects of **(b) illuminant condition** were significant for *optimal color model* and *mean LMS model* (*F*(1,9) = 12.5, *p* = 0.00636; *F*(1,9) = 8.78, *p* > 0.0159, respectively), but not significant for *mean chromaticity model* (*F*(1,9) = 4.81, *p* > 0.05). Again multiple comparison using a Bonferroni’s correction (significance level, 0.05) showed following results: (i) *optimal color model* > *mean LMS model*, *optimal color model* > *mean chromaticity model*, and *mean LMS model* > *mean chromaticity model* for *3000K*, and (ii) *optimal color model* > *mean LMS model*, *optimal color model* > *mean chromaticity model*, and *mean LMS model* = *mean chromaticity model* for *20000K*.

Overall, these statistical tests suggested that our optimal color model predicted observers’ settings generally better than the mean LMS and mean chromaticity models at least under the current experimental procedure. This observation was held for different distributions and test illuminants as supported by the results from multiple comparisons above.

Regarding the degree of color constancy, it was shown that human observers can achieve roughly 50% constancy for each distribution condition, though the *flat* condition showed slightly higher *CIs*. *MI*-based analysis revealed that our optimal color model predicted the shift of observers’ settings in response to test illuminant change better than other candidate models in general. With regard to raw chromaticity settings (Figure 5 (a)), our model was successful in explaining that there is little impact of the shape of distribution, but mean LMS model and mean chromaticity model also predicted this point (Figure 5 (b)). In Experiment 2, to further pursue the applicability of our hypothesis, we tested three different distributions designed to separate predictions made by our model and the mean LMS model. We designed the shape of color distributions so that optimal color model again perfectly predicts the chromaticity of illuminant but mean LMS model predicts a different illuminant for each distribution condition. Our interest was to see if the shape of the color distribution still has little effect on human observers’ white point settings as predicted by the optimal color model.

## 4. Experiment 2

The purpose of the Experiment 2 was to investigate human observers’ estimates of the illuminant for different distribution conditions, where the mean LMS model gave a biased estimation of the illuminant.

### 4.1 Color distribution of surrounding stimuli

In Experiment 2, we used three new distributions: *Red-reduced, Green-reduced* and *Blue-reduced*. We used the same 30 chromaticities as in Experiment 1, but luminance values were assigned differently as follows. First, as shown in Figure 11 (a), we divided the 30 chromaticities into 6 categories. There are chromaticities that are included in Red, Green and Blue groups only (shown by red, green and blue circles, respectively). Also, there are chromaticities that belong to more than one group (e.g. magenta circles belong to both Red and Blue groups). For *Red-reduced* distribution, colors that are included in the Red group (i.e. red, magenta and yellow symbols) were set to have one-third the luminance of optimal colors while other colors were all set to optimal colors. The same manipulation was applied to *Green-reduced* and *Blue-reduced* conditions. Our intension behind this manipulation is to reduce the contribution to mean LMS values from colors in a specific region in the chromaticity diagram. For example, for *Green-reduced* distribution colors that have low L/(L+M) values are forced to have low luminance, and thus the mean LMS values across 60 surfaces should be biased towards higher L/(L+M). We used 30 bright colors and 30 dark colors that have the same chromaticity but 20% of the luminance of the bright colors.

**Figure 11:**
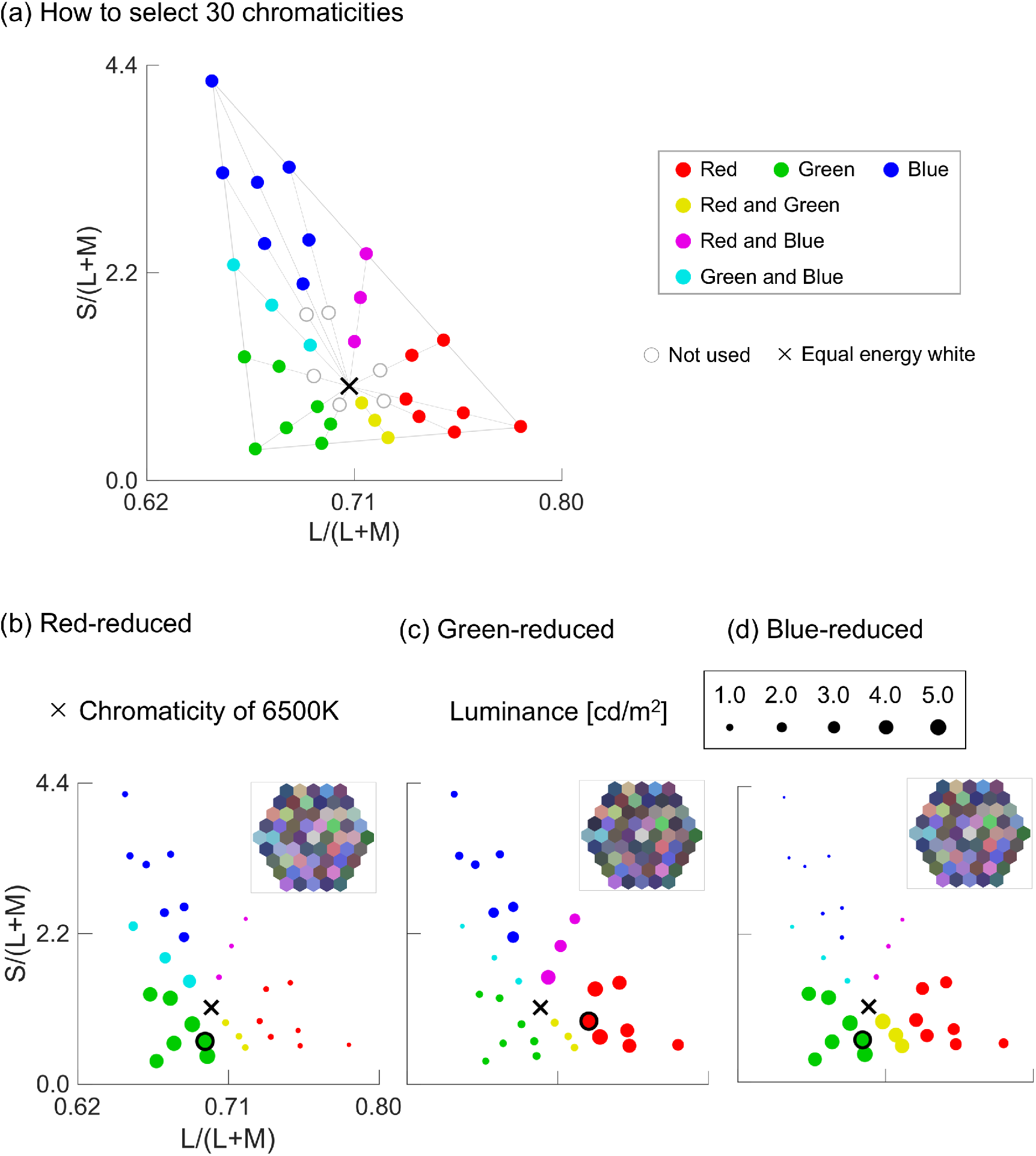
(a) How to group 30 chromaticities into six different categories. Note that colors marked by yellow, magenta and cyan symbols indicate that they belonged to both Red and Green, both Red and Blue, and both Green and Blue groups, respectively. (b) - (d) Color distributions for each distribution under *6500K*. The size of symbols expresses the luminance value at each chromaticity. The highest luminance color is marked by black edge. Only the 30 bright colors are shown here.

Figure 11 (b) - (d) show luminance values over chromaticities under the test illuminant of *6500K*. We see that for each distribution some colors have noticeably lower luminances than others. Inserted top-right figures indicate an experimental stimulus for each distribution condition, just to show how the impression of stimulus changes by manipulating the luminance distribution of surfaces without changing chromaticities.

In Experiment 2 we also set intensity of test illuminants such that mean luminance across the 60 colors became 1.2 cd/m^2^. To achieve this illuminant intensities were set to 4.45 cd/m^2^, 4.29 cd/m^2^, and 4.18 cd/m^2^ for *Red-reduced*, *Green-reduced*, and *Blue-reduced* in *3000K* condition. For *6500K*, they were 4.53 cd/m^2^, 4.63 cd/m^2^, and 4.63 cd/m^2^. Also, for *20000K*, we used 3.66 cd/m^2^, 3.74 cd/m^2^, and 3.80 cd/m^2^.

### 4.2 Results and discussion

The presentation of results follows the format in Experiment 1. Panel (a) in Figure 12 shows the averaged observer’s settings across 10 trials. The shape of symbol indicates the distribution condition (*Red-reduced*, *Green-reduced* and *Blue-reduced*), and colors indicate the illuminant condition (*3000K*, *6500K* and *20000K*).

**Figure 12:**
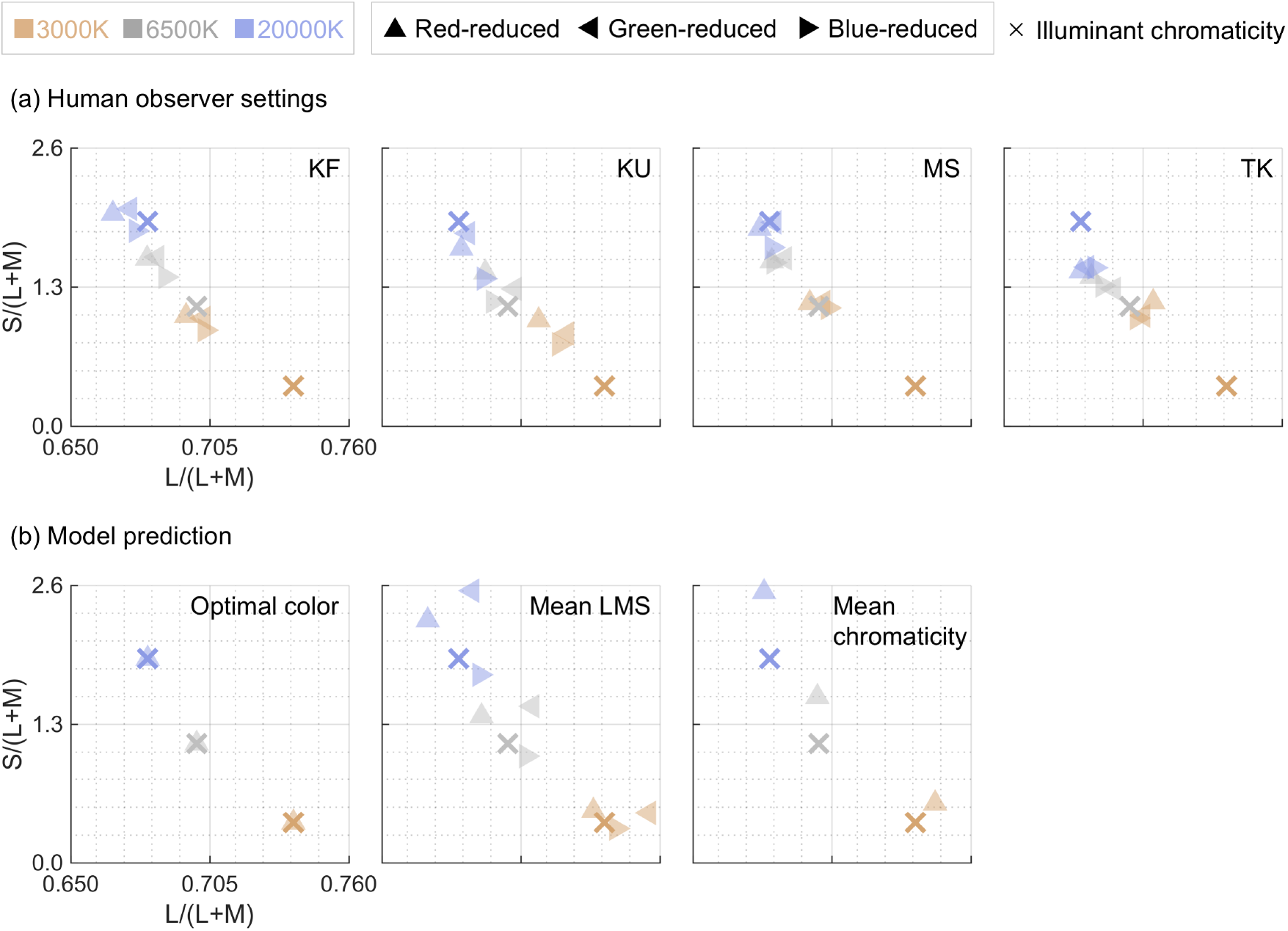
Observer’s white settings for each condition in Experiment 2. Different panels indicate different observers. (b) Model prediction by optimal color model, mean LMS model and mean chromaticity model. Optimal color model and mean chromaticity model predict the same chromaticity regardless of distribution condition by experimental design, and thus only prediction for *Red-reduced* condition is shown here.

Panel (b) shows predictions from optimal color model and mean LMS model, and mean chromaticity model. As in Experiment 1, optimal color model provided perfect estimation of illumination for all conditions by design, and thus we plotted only predictions for *Red-reduced* condition. Predictions from mean LMS show some differences depending on distribution condition as expected. For example, for *Red-reduced* condition, surfaces that have high L/(L+M) have lower luminance than other surfaces, giving less contribution to the averaged LMS values. As a result, the prediction shifted towards lower L/(L+M) direction. The mean chromaticity model predicted exactly the same chromaticity as Experiment 1 because our experimental manipulation changes only luminance profiles of surrounding stimuli while keeping chromaticities of 60 surfaces constant.

For observer MS and TK, we see that observers’ settings for three distribution conditions seem to come close for any illuminant condition, which is consistent with the prediction from optimal color model. However, some separation and systematic pattern of settings were observed for KU and especially KF: lower L/(L+M) for *Red-reduced* condition, higher L/(L+M) for *Green-reduced* condition, and lower S/(L+M) condition for *Blue-reduced* condition, which seems to be somewhat consistent with the predicted pattern by mean LMS model. Also, observer’s settings are predictably influenced by the color temperature of illuminant.

Figure 13 shows averaged luminance settings across 4 observers for each condition. We show the prediction from mean luminance, highest luminance and illuminant intensity by red, cyan and blue cross symbols, respectively. We performed two-way repeated-measures ANOVA for **(a) distribution condition** (*Red-reduced*, *Green-reduced* and *Blue-reduced*) and **(b) illuminant conditions** (*3000K*, *6500K* and *20000K*) as within-subject factors for the luminance settings. The main effect of **(a) distribution condition** and **(b) illuminant condition** were not significant (*F*(2,6) = 0.10, *p* > 0.05; *F*(2,6) = 1.65, *p* > 0.05, respectively). The interaction between two factors was not significant (*F*(4,12) = 0.55, *p* > 0.05), either. Thus, there was no significant difference for any pair of conditions in Experiment 2.

**Figure 13:**
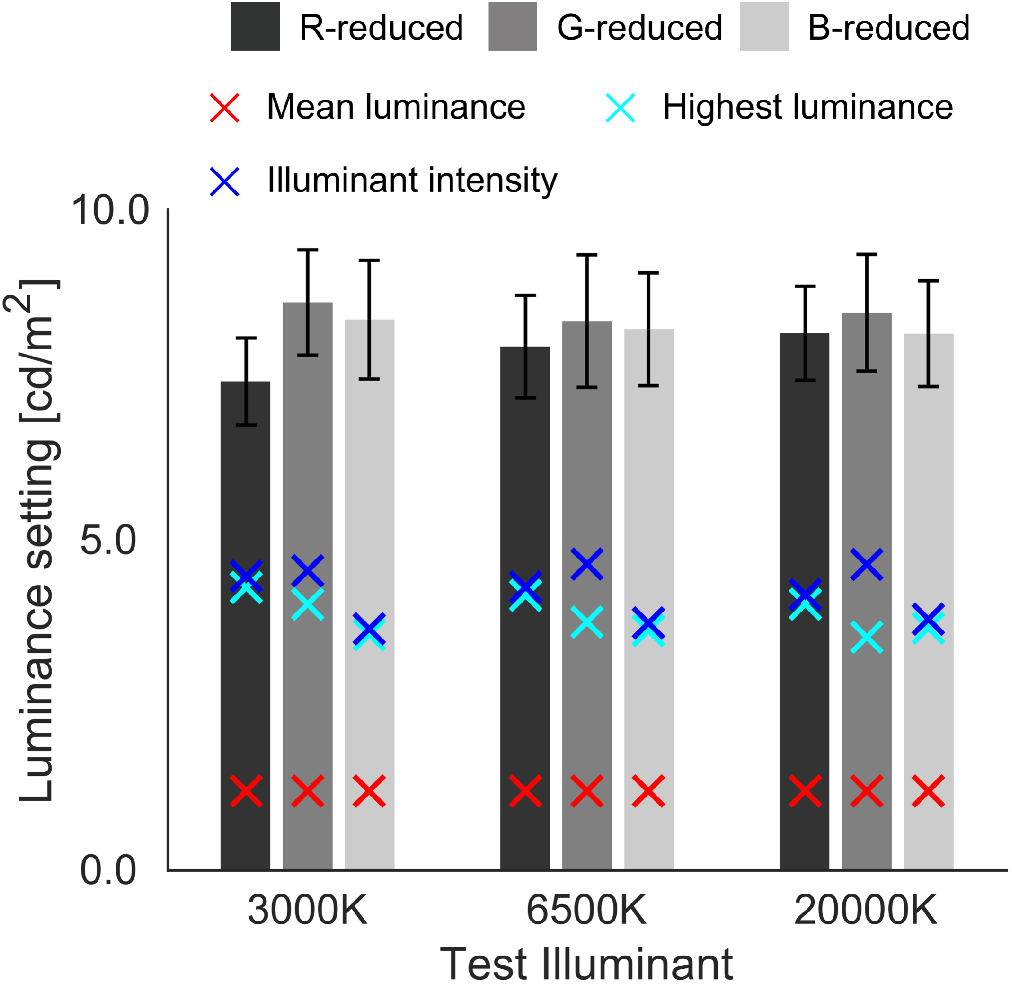
Observer’s luminance settings for each condition in Experiment 2. Distribution conditions are indicated by the lightness of the bars. Red and cyan crosses indicate the mean luminance and the highest luminance across 60 surrounding surfaces, respectively. Blue cross symbol shows the intensity of set test illuminant. Error bars are ± *S.E.* across four observers.

We again calculated correlation coefficient between nine observer’s settings and each luminance statistics. The correlation coefficients were −0.356 (*p* > 0.05), −0.6273 (*p* > 0.05) and 0.0210 (*p* > 0.05) for mean luminance, highest luminance, and illuminant intensity (i.e. prediction from optimal color model), respectively. Thus, these models failed to predict human observers’ performance. This might be due to the observers’ use of more complex strategy than simple luminance statistics tested here.

Next, we quantified the degree of color constancy based on equation (2). Figure 14 shows the *CIs* for all conditions. We conducted a two-way ANOVA for **(a) distribution condition** (*Red-reduced*, *Green-reduced* and *Blue-reduced*), and **(b) illuminant conditions** (*3000K* and *20000K*) as within-subject factors for the *CIs*. The main effect of **(a) distribution condition** was not significant (*F*(2,6) = 3.59, *p* > 0.05). Also, the main effect of **(b) illuminant condition** and the interaction between two factors were not significant (*F*(1,3) = 7.18, *p* > 0.05; *F*(2,6) = 2.16, *p* > 0.05, respectively). Consequently, the degree of color constancy was not influenced by the shape of distribution nor the color temperature of test illuminants.

**Figure 14:**
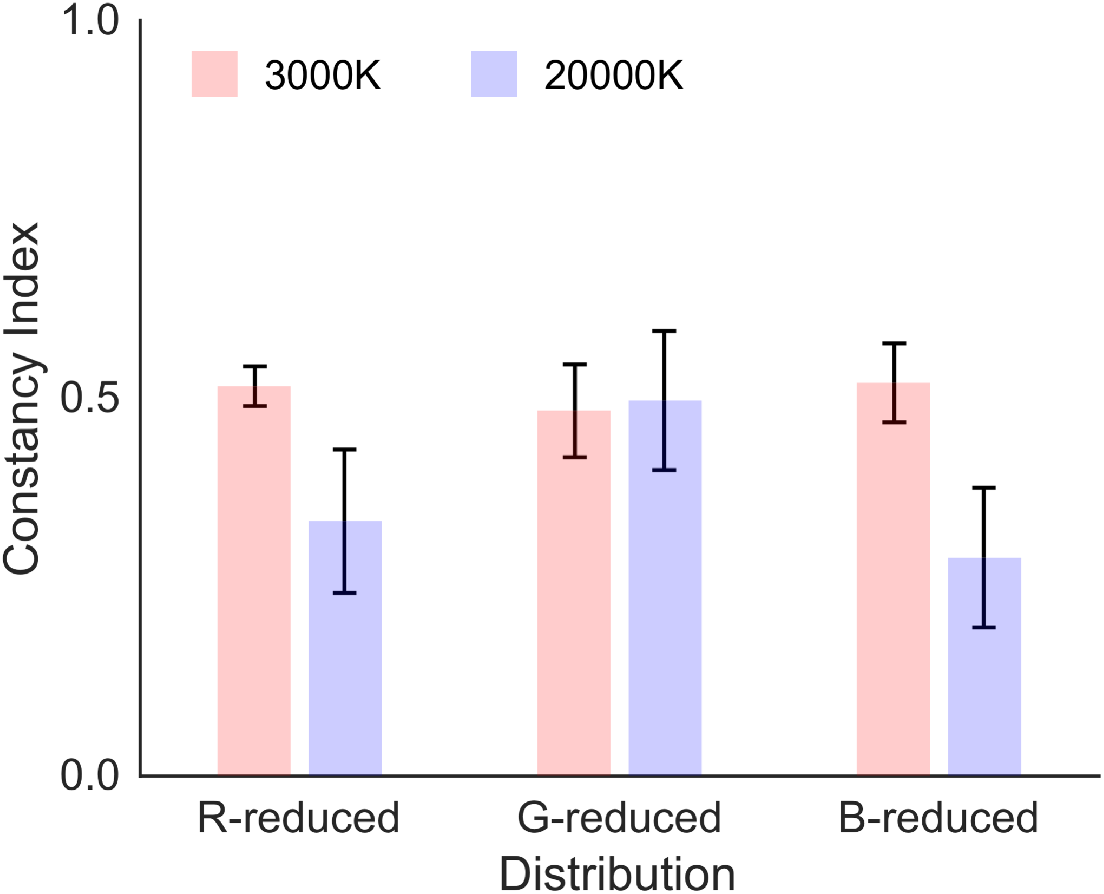
Constancy index (*CI*) for each condition in Experiment 2. *CIs* closer to 1.0 indicates higher degree of color constancy. Error bars show ±*S.E.* across 4 observers.

We again calculated *MIs* from equation (3) based on the prediction from optimal color, mean LMS and mean chromaticity models as shown in Figure 15. For *3000K* condition (left 9 bars), it was shown that the values of *MI* are generally higher for optimal color model than other models, except for *Blue-reduced* condition where mean LMS shows a slightly higher *MI* than that of optimal color model. This observation holds also for *20000K* condition. However, error bars are found to be large, making it hard to make a conclusive statement whether our optimal color model overall worked better than other models. Thus, we performed a following statistical analysis.

**Figure 15:**
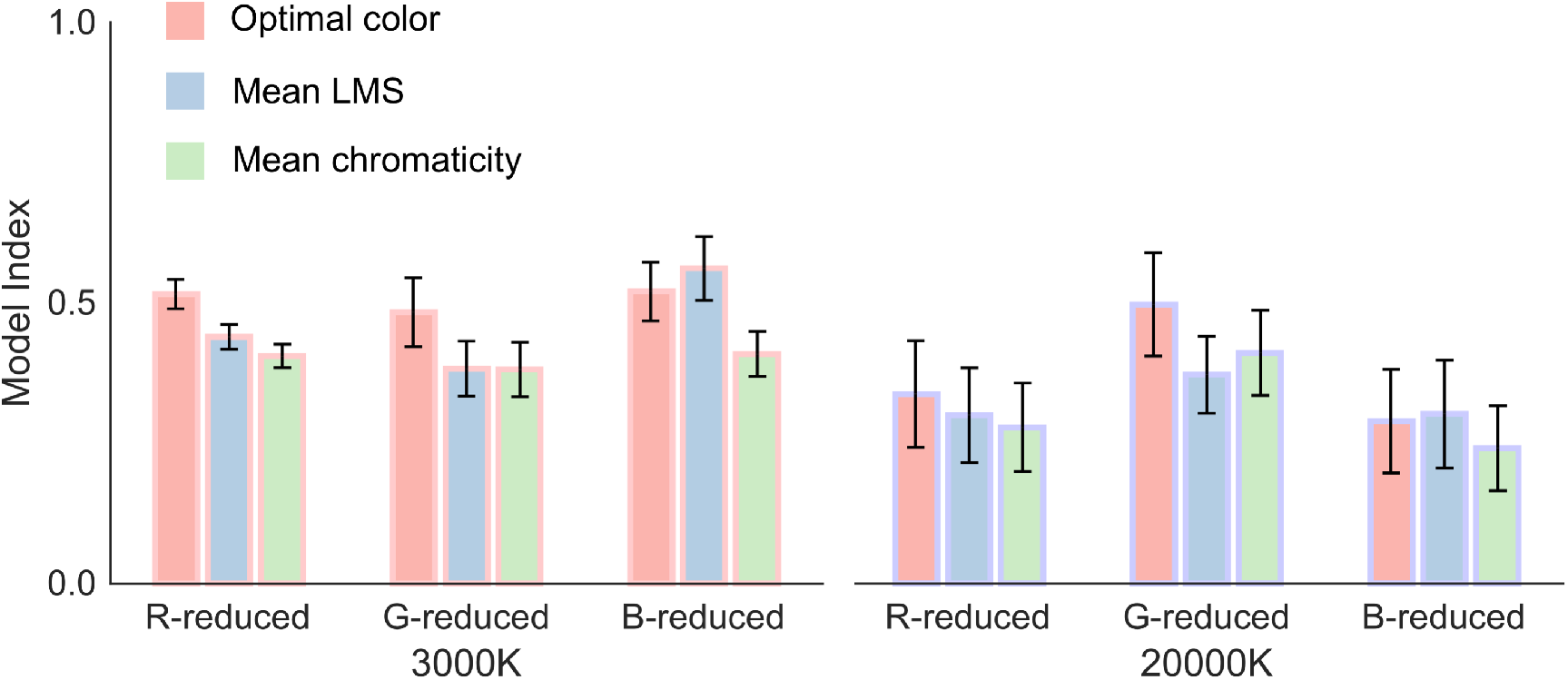
Model index (*MI*) for each condition in Experiment 2. Higher values indicate better prediction by models. Error bars show ±*S.E.* across 4 observers.

A three-way repeated-measures analysis of variance (ANOVA) were conducted for **(a) model types** (*optimal color, mean LMS* and *mean chromaticity*), **(b) distribution condition** (*Red-reduced*, *Green-reduced* and *Blue-reduced*), and **(c) illuminant condition** (*3000K* and *20000K*) as within-subject factors for the *MIs*. The main effect of **(a) model type** was significant (*F*(2,6) = 68.1, *p* = 0.000075) while the main effect of **(b) distribution condition** and **(c) illuminant condition** were not significant (*F*(2,6) = 1.27, *p* > 0.05; *F*(1,3) = 6.82, *p* > 0.05). The interaction between **(a) model type** and **(b) distribution condition** and between **(a) model type** and **(c) illuminant condition** were significant (*F*(4,12) = 48.1, *p* < 0.00001; *F*(2,6) = 18.6, *p* < 0.00268, respectively). In contrast, the interaction between **(b) distribution condition** and **(c) illuminant condition** was not significant (*F*(2,6) = 2.18, *p* > 0.05). The interaction among three factors was not significant, either (*F*(4,12) = 3.02, *p* > 0.05).

Regarding the significant interaction between **(a) model type** and **(b) distribution condition**, the simple main effect of **(a) model type** was all significant at *Red-reduced*, *Green-reduced* and *Blue-reduced* conditions (*F*(2,18) = 38.7, *p* < 0.00001; *F*(2,18) = 75.6, *p* < 0.00001; *F*(2,18) = 64.0, *p* < 0.00001, respectively). In addition, the simple main effect of **(b) distribution condition** was significant for optimal color model (*F*(2,18) = 4.79, *p* = 0.0184), but not significant for mean LMS model and mean chromaticity model (*F*(2,18) = 2.79, *p* > 0.05; *F*(2,18) = 3.36, *p* > 0.05). Furthermore, multiple comparison using a Bonferroni’s correction (significance level, 0.05) showed a following relation: (i) *Red-reduced* = *Green-reduced*, *Red-reduced* = *Blue-reduced*, and *Green-reduced* > *Blue-reduced* for *optimal color model*. Also, a following result was observed: *optimal color* > *mean LMS*, *optimal color* > *mean chromaticity*, *mean LMS* > *mean chromaticity* for *Red-reduced* and *Blue-reduced* conditions, and *optimal color* > *mean LMS model*, *optimal color* > *mean chromaticity*, *mean LMS model* = *mean chromaticity* for *Green-reduced* condition.

Next, we analyzed simple main effects of interaction between **(a) model type** and **(c) illuminant condition**. We found that **(a) model type** showed significant simple main effects at *3000K* and *20000K* (*F*(2,12) = 79.7, *p* < 0.00001; *F*(2,6) = 31.0, *p* = 0.000018, respectively). Moreover, the simple main effects of **(b) illuminant condition** were significant for *optimal color model* and *mean LMS model* (*F*(1,9) = 8.27, *p* = 0.0183; *F*(1,9) = 8.88, *p* = 0.0155, respectively), but not significant for *mean chromaticity model* (*F*(1,9) = 3.73, *p* > 0.05). Again, multiple comparison using a Bonferroni’s correction (significance level, 0.05) showed a following result: (i) *optimal color* > *mean LMS*, *optimal color* > *mean chromaticity*, *mean LMS* > *mean chromaticity* at *3000K*, and (ii) *optimal color* > *mean LMS*, *optimal color* > *mean chromaticity*, *mean LMS* = *mean chromaticity* at *20000K*.

Our main interest in this experiment was to evaluate whether our proposed model preforms better than other candidate models. In summary, the calculation of *MIs* and these statistical analyses overall supported this idea that our proposed model accounts for observers’ settings better than the mean LMS and the mean chromaticity models, which is consistent with the trends found in Experiment 1. This claim seems to generally hold for the tested distribution shape in Experiment 2.

However, the prediction from optimal color model is still limited in Experiments 1 and 2 as *MIs* for the optimal color model were considerably smaller than 1.0, showing the lack of agreements. Thus, it is still inconclusive as to whether the optimal color model can be a good candidate strategy adopted by human observers to infer the influence of illuminant.

Regarding a limited account of our model, it is worth noting that we used optimal surfaces for surrounding stimuli which do not exist in a real world, and thus observers never experienced luminance distributions formed by optimal colors. Instead, in the real world, the visual system encounters a scene containing natural objects that have a reflectance less than 1.0; therefore human color constancy may work better in such a condition. Also, such a natural scene is more difficult for the model as the model also needs to guess the influence of the illuminant, which will lead to an uncertainty of inference and the model presumably begins to make mistakes. Therefore, we decided to design scenes with a color distribution formed by natural objects and to test our hypothesis in a set-up which is challenging to both human observers and the proposed model.

In the following Experiment 3, we used 60 reflectances selected from 575 natural object database (Brown, 2003). We manipulated the shape of distribution to “deceive” the model so that even when the test illuminant is held constant, the model predicts different illuminant chromaticities influenced by the shape of the color distribution. Then we measured observers’ white points using the same procedure as in Experiment 1 and 2. We tested whether (i) human observers’ white points also change in response to the change of the shape of color distribution, and if so whether the (ii) optimal color model can predict those shift patterns in observer’s white points.

## 5. Experiment 3

### 5.1 Color distribution of surrounding stimuli

We used three distributions: *natural, Red-increased and Blue-increased*. As argued in the previous section, the aim of manipulating color distribution in this experiment was to alter the prediction by the optimal color model. To achieve this, we selected 60 reflectances from a database that contains 575 spectral reflectances of natural objects (Brown, 2003) based on the following criteria. First, since a white surface provides a direct cue to the illuminant color, it allows observers to adjust the color of the test field so that it simply matches the white surface. Therefore, we decided to exclude 59 flat reflectances that have chromaticities in a range between 0.6978 and 0.7178 along L/(L+M) axis and between 0.800 and 1.200 along S/(L+M) axis when rendered under equal energy white. Secondly, we randomly sampled 60 surfaces from the remaining 516 reflectances so that mean chromaticity across the 60 surfaces corresponds to the chromaticity of equal energy white (0.7078 and 1.0000 for L/(L+M) and S/(L+M), respectively) when rendered under equal energy to make a scene chromatically balanced. The chosen 60 surfaces were defined as the *natural* distribution. We also made sure that for the *natural* distribution condition the optimal color model can perfectly estimate the chromaticity of the test illuminant.

Next we manipulated the luminance profile of *natural* distribution to alter the model prediction without changing chromaticities in the following ways. To create the *Red-increased* condition, we scaled up or down 60 reflectances by multiplying scalar values without changing the spectral shape of the reflectances. The scalar value was determined proportionally to the L/(L+M) of the reflectance under equal energy white so that reflectances with higher L/(L+M) have higher luminance. Similarly, to create the *Blue-increased* distribution, we scaled 60 reflectances by the scalar value as a function of S/(L+M) so that the reflectances with high S/(L+M) gets lighter. We made sure that maximum reflectance does not exceed 1.0 to keep all reflectances physically plausible.

Resultant distributions are shown in Figure 16. We see that the (a) *natural* distribution has a mountain-like shape, whilst the (b) *Red-increased* and the (c) *Blue-increased* conditions have skewed luminance profiles. For the (b) *Red-increased* distribution, the upper-subpanel shows that luminances around higher L/(L+M) regions are elevated. Also, for the (c) *Blue-increased* distribution, the lower-subpanel shows that luminances around higher S/(L+M) regions are increased. Importantly, fitted optimal color distributions are plotted together, and best-fit color temperature and illuminant intensity are shown at the top-right of each upper sub-panel. These fitting results indicate that model perfectly predicts the color temperature for the (a) *natural* condition. In contrast, the estimation is biased in 5500K and 8000K for the (b) *Red-increased* and the (b) *Blue-increased* distributions, respectively, as expected. This was also the case for *6500K*. For the *4000K* condition, the model prediction of color temperature was 4000K, 3500K, and 4500K for the *natural*, *Red-increased*, and *Blue-increased* distributions. For the *10000K* condition, the prediction was 10000K, 7500K, and 15500K.

**Figure 16:**
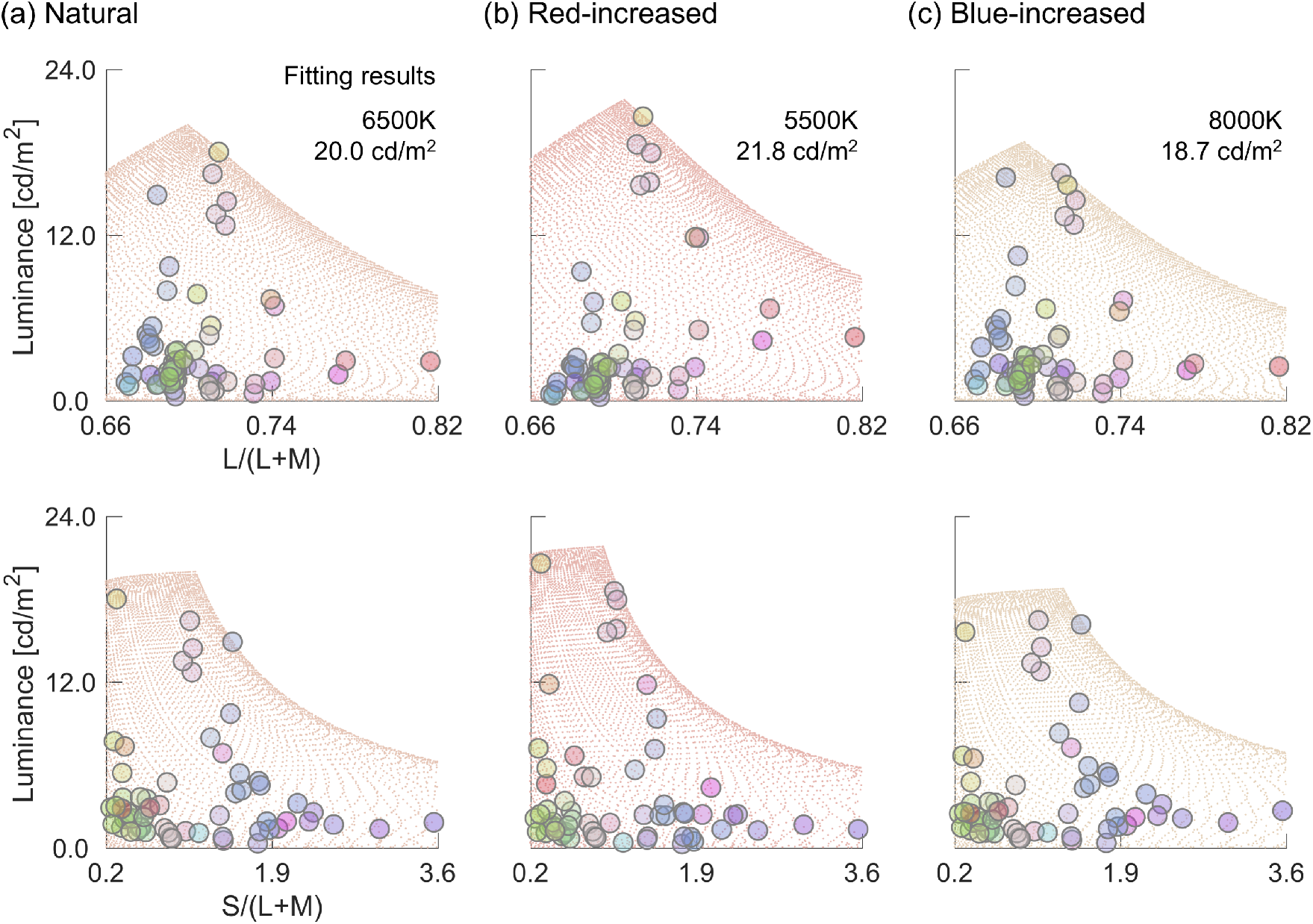
Color distributions for each distribution condition under *6500K* in Experiment 3. Each panel also indicates the optimal color distribution best-fit to the 60 colors.

Note that different color temperatures were used for test illuminants in Experiment 3 (*4000K*, *6500K* and *10000K*) compared to Experiments 1 and 2 (*3000K*, *6500K* and *20000K*). This is because we wanted to manipulate distribution shapes so that the model predicts higher or lower color temperature than the ground-truth. If we instead use 20000K as a test illuminant, the model might predict a color temperature 30000K for the *Blue-increased* distribution, and observers’ chromaticity settings might exceed the chromatic gamut of the CRT display. For this reason, we employed less extreme color temperatures in this experiment.

We set intensities of test illuminants so that the mean luminance across 60 surfaces becomes 6.0 cd/m^2^ for all conditions. To achieve this, illuminant intensities were set to 25.1 cd/m^2^, 25.4 cd/m^2^, and 25.6 cd/m^2^ for *natural*, *Red-increased*, and *Blue-increased* distributions in *4000K* condition. For *6500K*, they were 26.3 cd/m^2^, 26.9 cd/m^2^, and 27.3 cd/m^2^. For *10000K*, we used 27.4 cd/m^2^, 27.7 cd/m^2^, and 27.8 cd/m^2^.

### 5.2 Results and discussion

Figure 17 shows the average of human observers’ settings across 20 settings for each condition. The presentation of results follows previous experiments.

**Figure 17:**
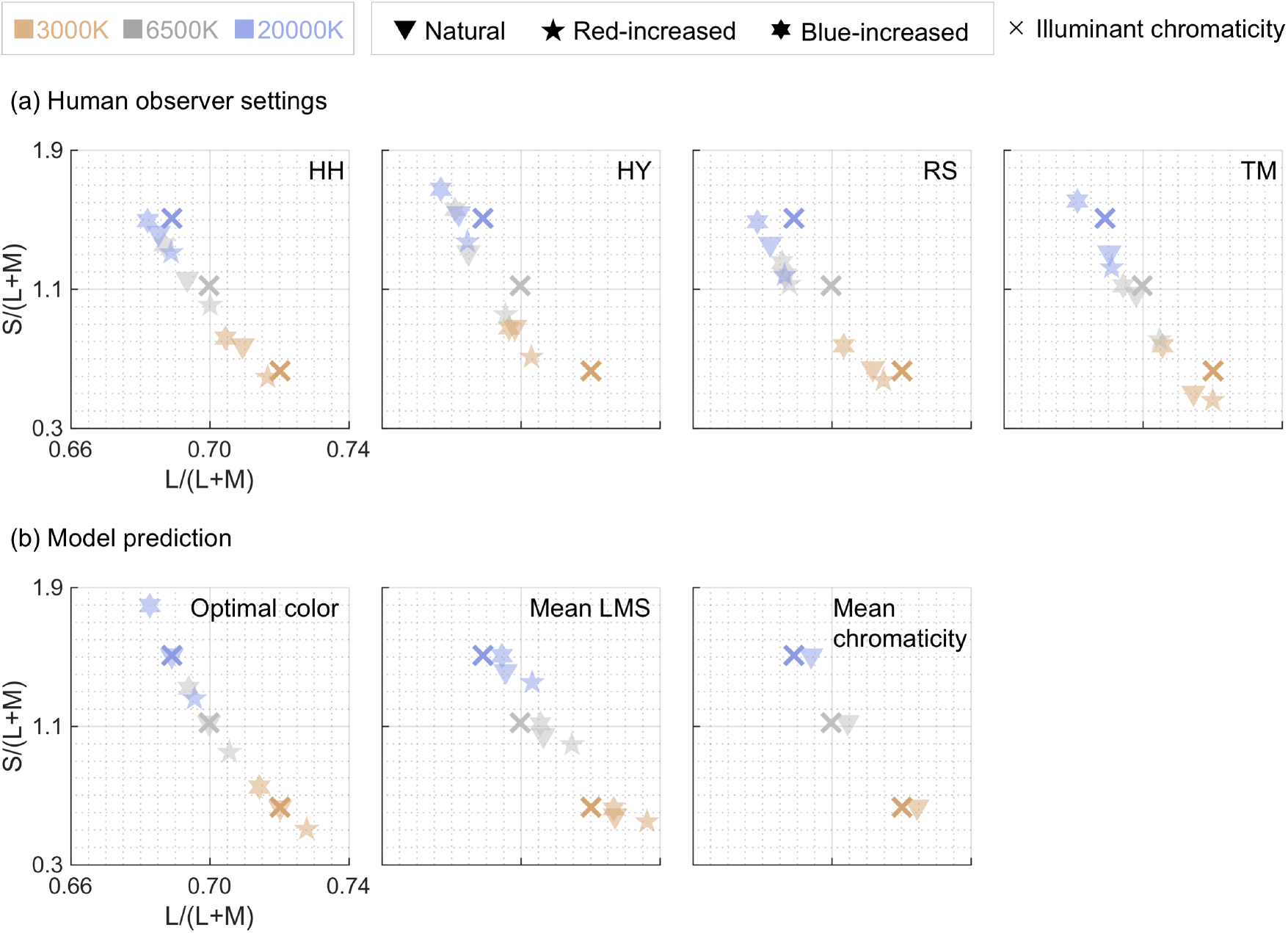
Observer’s chromatic settings for each condition in Experiment 3. Different panel indicates a different observer. (b) Model prediction by optimal color model, mean LMS model and mean chromaticity model. Mean chromaticity model predicts the same chromaticity for any distribution condition by design, and thus only prediction for *Natural* condition is shown here.

In this experiment, the panel (b) is particularly important as the optimal color model provided different chromaticities for each distribution condition by design. We here see that the prediction by mean LMS model is also influenced by the shape of distribution. Mean chromaticity model predicted the same chromaticity for any distribution condition, and thus any shift of observers’ perceptual white point depending on distribution condition cannot be explained by mean chromaticity model.

The panel (a) shows that all observers’ settings exhibit a systematic pattern as follows. When the distribution is *Red-increased*, observers’ settings shifted towards higher L/(L+M) and lower S/(L+M) direction compared to settings under *natural* distribution. When the distribution is *Blue-increased*, observers settings shifted towards lower L/(L+M) and higher S/(L+M) direction. Thus, the shape of color distribution has strong effects on observers’ settings in this experiment. Importantly these shifts closely resemble the prediction pattern by optimal color model as shown in panel (b). These results would support our hypothesis that human observers use the shape of color distribution to infer the chromaticity of illuminant. However, note that mean LMS model seems to predict the shift of observer’s settings reasonably well, too. Later in this section, we quantify the degree of agreement between observers’ settings and model prediction based on correlation coefficient and *RMSE* values.

Figure 18 shows the luminance settings for each condition. Overall, it is shown that observers’ luminance settings are closer to the illuminant intensity (i.e. ground-truth) compared to Experiments 1 and 2 (see Figure 6 and 13), suggesting better estimation of the intensity of illuminants. We performed two-way repeated-measures ANOVA for **(a) distribution condition** (*natural, Red-increased, and Blue-increased*), and **(b) illuminant conditions** (*4000K*, *6500K* and *10000K*) as within-subject factors for the luminance settings. The main effects of **(a) distribution condition** and **(b) illuminant condition** were not significant (*F*(2,6) = 0.149, *p* > 0.05; *F*(2,6) = 4.06, *p* > 0.05, respectively). The interaction between two factors was not significant (*F*(4,12) = 1.32, *p* > 0.05), either. Thus, we found no significant difference for any pair of conditions in Experiment 3.

**Figure 18:**
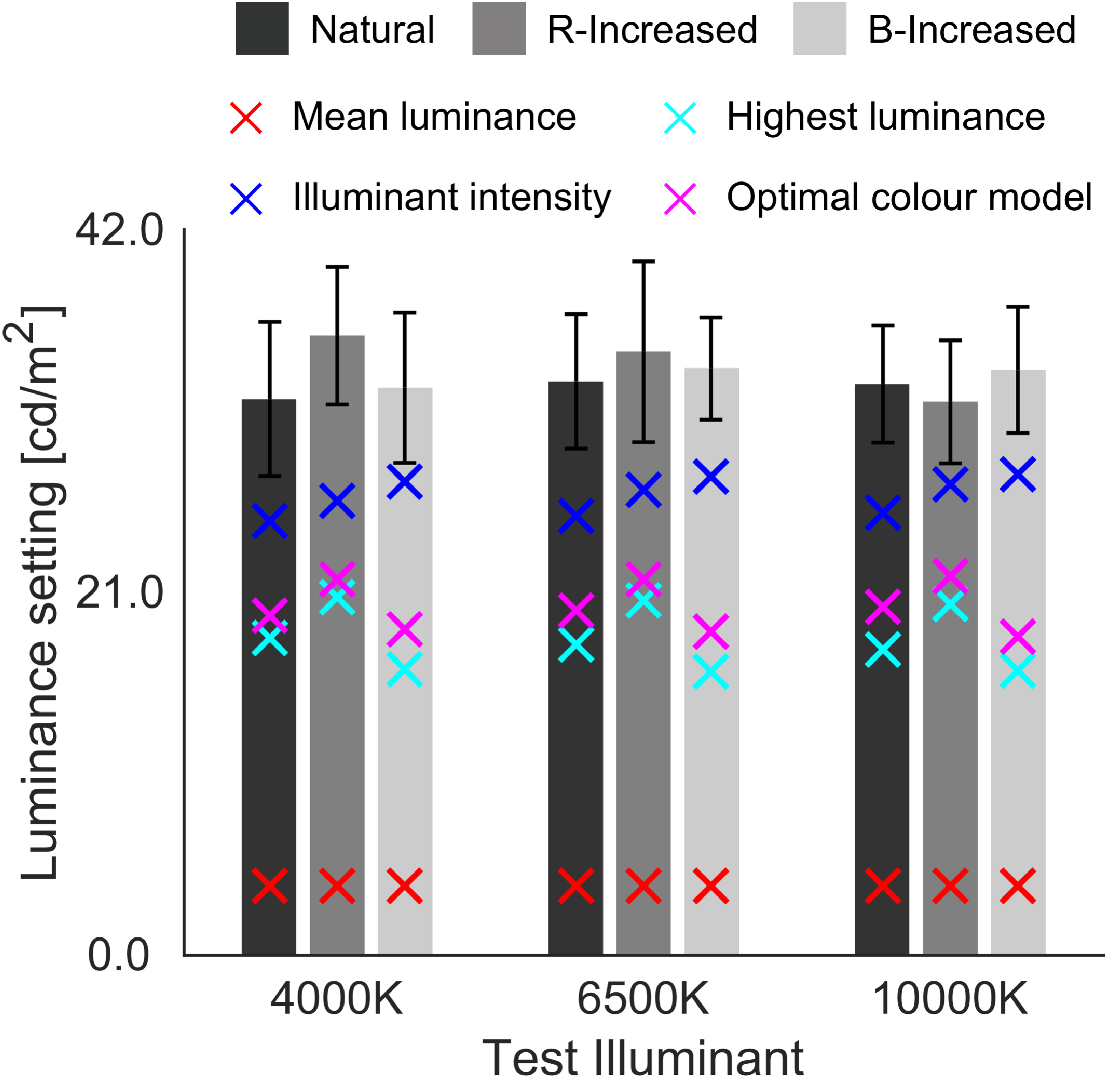
Observer’s luminance settings for each condition in Experiment 3. Red and cyan crosses indicate the mean luminance and the highest luminance across 60 surrounding surfaces, respectively. Blue cross symbol shows the intensity of set test illuminant. The magenta cross denotes the prediction by optimal color model. Error bars are ± *S.E.* across four observers.

We again calculated correlation coefficient between averaged observer’s settings (9 points) and each luminance statistics. Correlation coefficients were 0.2154 (*p* > 0.05), 0.3255 (*p* > 0.05), 0.1833 (*p* > 0.05) and 0.2701 (*p* > 0.05) for mean luminance, highest luminance, illuminant intensity and optimal color model, respectively. Thus, any of the tested models here does not show a particularly high correlation. This result is in agreement with Experiment 2.

Since the main focus in this experiment is to examine the effect of distribution shape on observers’ settings, we do not show *CIs* which focuses on the effect of illuminant change on observers’ settings. Instead, we argue the degree of agreement between model prediction and observers’ settings using correlation coefficients. Figure 19 shows the plot where horizontal axis is the prediction of optimal color model and vertical axis is the observers’ settings for (a) L/(L+M) and (b) S/(L+M) directions. The dotted line indicates a unity line. The solid line is a straight line fitted to the nine data points. The correlation coefficient is shown at the bottom-right in each sub-panel where *** indicates *p*-value less than 0.001. Correlation coefficient shows fairly high values, suggesting that optimal color model can account for human observer’s behavior well.

**Figure 19:**
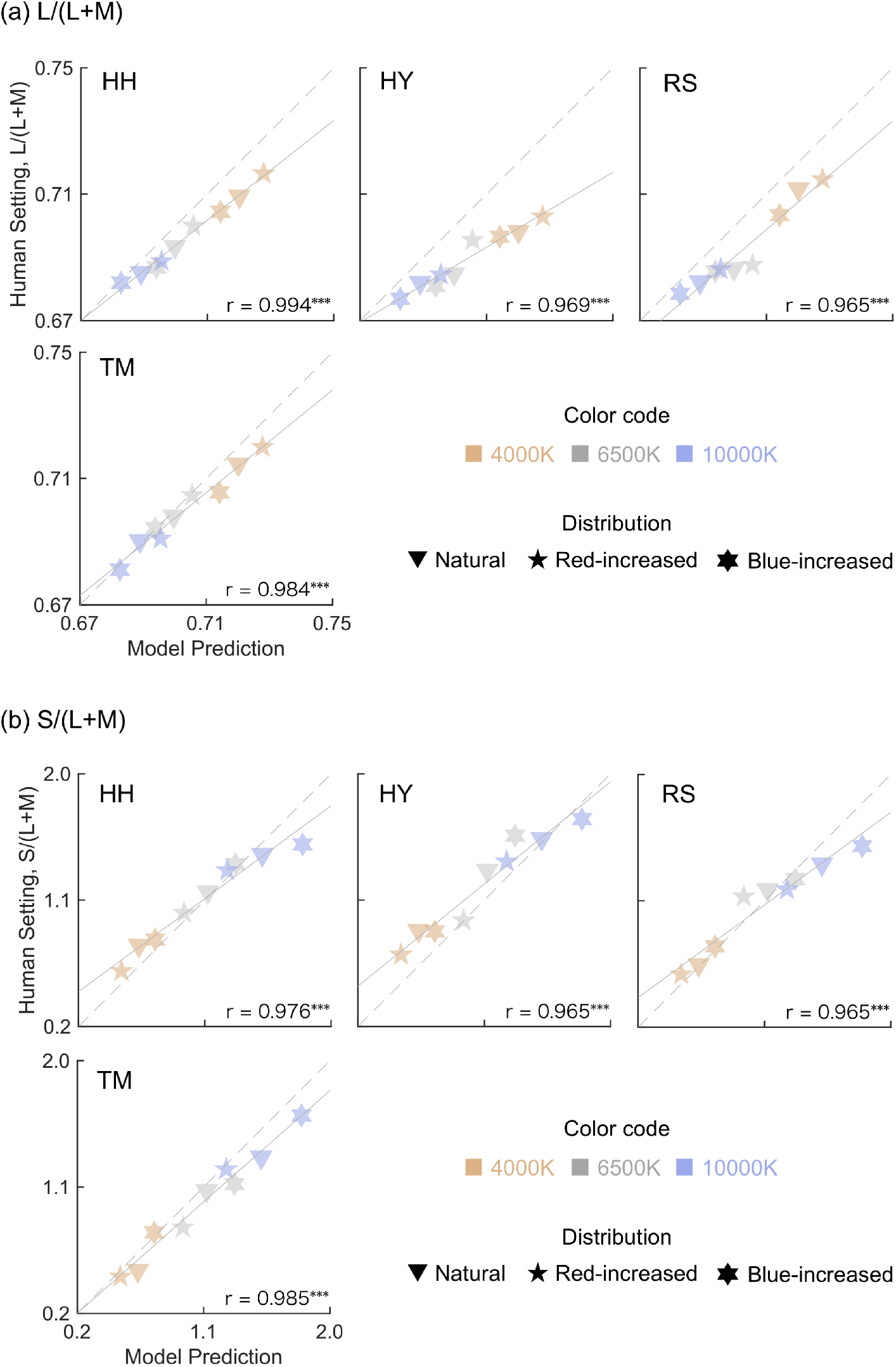
(a) Optimal color model prediction (horizontal axis) vs. human observers’ settings (vertical axis) for L/(L+M) direction. Different sub-panels indicate different observers. The right-bottom numbers show correlation coefficient and *** marks indicate p-value less than 0.001. The dotted straight line shows a unity line and solid line shows straight line fit to the data points. (b) The same plot for S/(L+M) direction.

However, whilst these assessments allow us to see the degree of correlation between model prediction and observers’ settings, data points should all fall on the unity line if a model prediction is perfect. For example, we see that data points are generally slightly off downward from the unity line for (a) L/(L+M) direction, especially for 4000K condition. To quantify the deviation of data points from the unity line, we calculated the *RMSE* values between model prediction and observer settings, across 9 data points separately for L/(L+M), S/(L+M) directions. Correlation coefficient and *RMSE* together should allow us to conclude the accuracy of model prediction.

In Figure 20, we summarize correlation coefficient (upper sub-panels) and *RMSE* values (lower sub-panels) for optimal color model and other models. We calculated both indices separately for each observer first (as demonstrated in Figure 19), and Figure 20 shows the averaged values across 4 observers.

**Figure 20:**
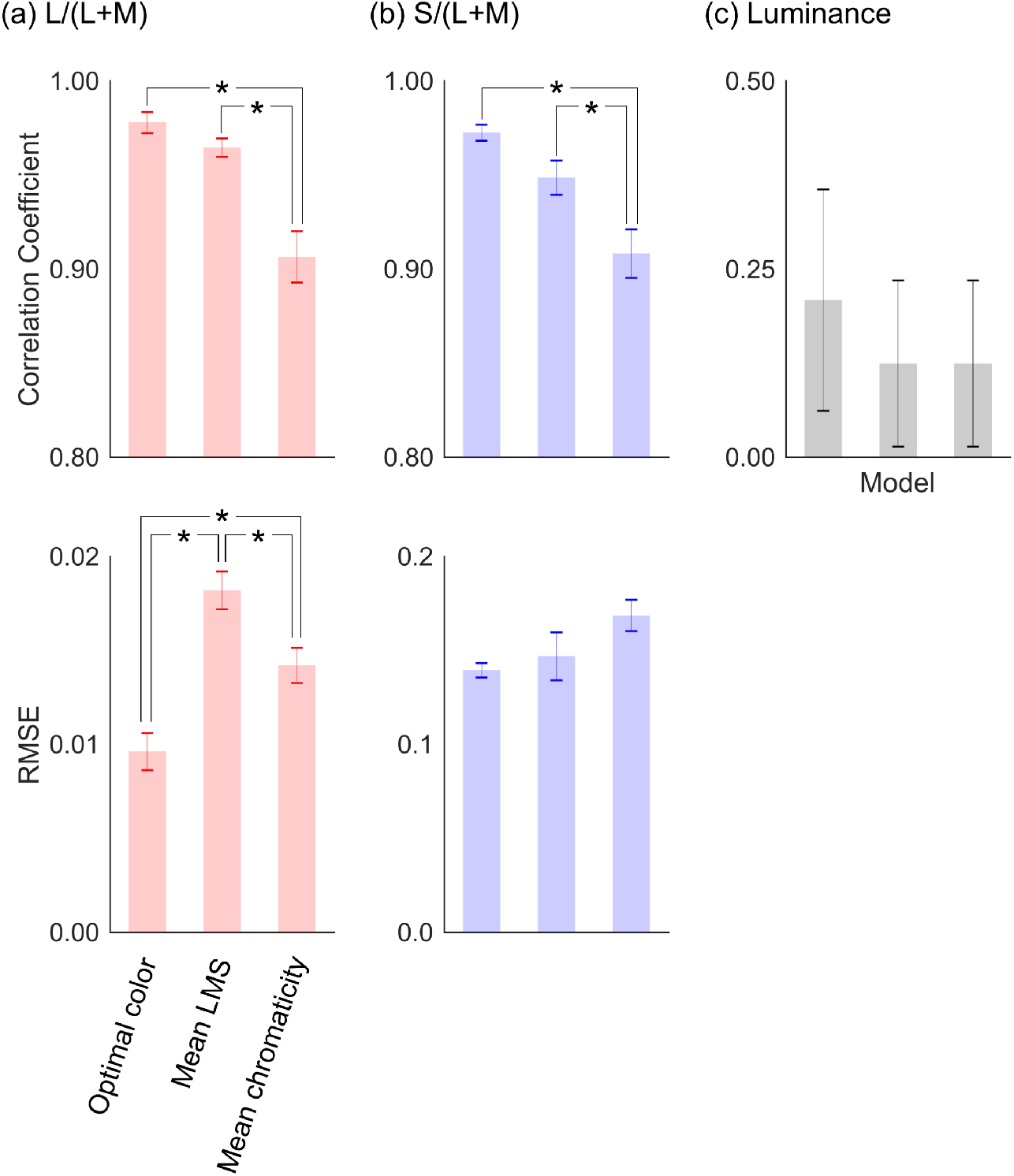
Correlation coefficient (upper panel) and RMSE (lower panel) between observer’s settings and model prediction for (a) L/(L+M), (b) S/(L+M) and (c) Luminance. Values are averaged across 4 observers, and error bars indicate ± *S.E.* across four observers. Note that the range of vertical axis is different for (c) Luminance direction. Asterisks * denote a significant difference (α < 0.05, Bonferroni’s correction).

Here, upper sub-panels show that optimal color model indicates the highest correlation coefficient for both chromatic channels. Mean LMS shows a correlation close to optimal color model. Mind that the range of vertical axis is between 0.80 and 1.00 for (a) L/(L+M) and (b) S/(L+M) indicating very high correlations for any tested model here. For (c) luminance, the correlation coefficient is noticeably lower than (a) L/(L+M) and (b) S/(L+M) for any model. Also, variation among observers seem to be larger as indicated by the length of error bars. Mean LMS and mean chromaticity model give the same estimation of illuminant intensity, and thus the correlation coefficients are also exactly the same. Regarding *RMSE* values, optimal color model shows a lower value for (a) L/(L+M) than other two models. Also, the trend is held for (b) S/(L+M) direction, but the differences across models seem to be small. We do not show *RMSE* values for (c) luminance channel, as we believe that the absolute prediction from mean LMS and mean chromaticity models should not match with human observers’ luminance settings and that only a relative measurement matters here (e.g. correlation coefficient).

For correlation coefficient, we performed one-way repeated-measures ANOVA, separately for each channel ((a) *L/(L+M)*, (b) *S/(L+M)* and (c) *luminance*), for model type (*optimal color*, *mean LMS*, and *mean chromaticity models*) as within-subject factor. The main effect of the model type was significant for (a) L/(L+M) and (b) S/(L+M) (*F*(2,6) = 15.8, *p* = .00408; *F*(2,6) = 21.0, *p* = 0.00195, respectively), but not for (c) luminance (*F*(2,6) = 0.0094, *p* > 0.05). We next performed multiple comparison using Bonferroni’s correction (significance level α = 0.05). We show pairs that showed significant differences by asterisks (*) in Figure 20.

Also, for *RMSE* values, one-way repeated-measures ANOVA was conducted separately for (a) *L/(L+M)*, *(b) S/(L+M)* and (c) *luminance* channels, for model type (*optimal color*, *mean LMS*, and *mean chromaticity*) as within-subject factor. The main effect of the model type was significant for (a) *L/(L+M)* and (c) *luminance* (*F*(2,6) = 1687.0, *p* < 0.00001; *F*(2,6) = 7.41 *p* = 0.0240, respectively), but not for (b) *S/(L+M)* (*F*(2,6) = 1.44 *p* > 0.05). We next performed multiple comparison using Bonferroni’s correction (significance level α = 0.05), and pairs that showed a significant difference are highlighted by asterisks in Figure 20.

Overall, these results show that the optimal color model is better than the mean LMS model and the mean chromaticity model in predicting observers’ estimations of L/(L+M) and that tested models are equally good at predicting observers’ estimation of S/(L+M). Also, in terms of predicting intensity estimation, none of tested models here was particularly good.

In summary of Experiment 3, a proposed optimal color model accounted for the shift of chromatic settings in response to the change of the scene color distribution with high precision. Here we note that our experiment still should not rule out the possibility of visual system’s use of the mean LMS model or any other models that we did not test in the present study. Nevertheless, empirical data here suggests that this geometry-based estimation of illumination might be a plausible algorithm for our visual system to infer the influence of illumination on a scene.

## 6. General discussion

From the lights reflected back from surfaces, how can we infer the color of the illuminants projecting onto the surface? This is the question that our visual system faces when solving color constancy. An intuitive answer would be to use the chromaticities of surfaces (as for example when the illuminant is reddish, chromaticities of all surface gets reddish). What is less known is that the luminance distribution also systematically changes, providing a diagnostic cue to the illuminant color. Our present study concerns this importance of the luminance distribution and specifically investigated whether subjective white point settings recorded from human observers can be predicted by the optimal color model, which makes an attempt to estimate illuminant color based on the shape of the color distribution. We conducted three psychophysical experiments to test this hypothesis. Experiment 1 and 2 were designed so that the optimal color model predicts a constant illuminant despite the change of distribution shapes and observers’ behavior generally agreed with this model prediction. In Experiment 3 we manipulated the shape of color distribution so that the optimal color model predicts the *success* or *failure* of color constancy and we found that observers’ chromaticity settings followed the prediction. These empirical data collectively suggest that the optimal color model could be a good candidate model for human color constancy, especially in estimating the chromaticity of illuminants in scenes that contain natural objects.

However, the model account regarding observers’ illuminant intensity estimation was largely limited. One notable feature in results was that observers’ luminance settings were substantially higher than actual illuminant intensity (shown in Figures 6, 13, and 18). This implies that human observers assumed that presented surrounding stimuli are darker than optimal colors because under the estimated illuminant intensities presented stimuli are not optimal colors. If we think that optimal colors do not exist in a real world, it makes sense that observers assumed the high intensity of the illuminant rather than high surface spectral reflectance. In the framework of the proposed model, there is strong association between accuracies in illuminant intensity estimation and in color temperature estimation. To demonstrate this let us consider a scene illuminated by a bluish illuminant. Under the bluish illuminant bluish surfaces should get lighter and thus one would expect that optimal color distribution under a reddish illuminant would be inappropriate, as it cannot cover light blue surfaces. However, this is true only when the illuminant intensity is properly estimated. In theory, if we assume a very intense illuminant, the optimal color distribution of any color temperature should cover the blue surface. In other words, for the model to estimate the color temperature precisely the illuminant intensity must be estimated well too. Thus, inaccurate illuminant intensity estimation by human observers could be one reason for imperfect agreement between model prediction and observer settings. This conclusion is also supported by the result that observers’ estimations of illuminant intensity was better in Experiment 3.

It should be noted that the present study used flat and matte experimental stimuli that are essentially an array of colors, to eliminate other cues to the illuminant, for example the influence of memory color (Olkkonen, Hansen and Gegenfurtner, 2008). In contrast, objects in real world are usually three-dimensional and sometimes contain specular reflection. Thus a single object typically exhibits a color variation over a surface which may provide additional cues to color constancy. The use of simple stimuli may be one of the potential reasons why we found a relatively low degree of color constancy of around 50% in Experiments 1 and 2. Especially the use of three-dimensional stimuli (Morimoto et al., 2017; Mizokami, 2019) and the presence of specular reflection (Yang and Shevell, 2012; Lee, and Smithson, 2016) are claimed to be important for color constancy. Thus, further extension of the present study using more realistic experimental stimuli would be desired. However, to put it another way, our experimental set-up was rather difficult situation for human observers compared to richness of illuminant cues in the real world, and it may be remarkable that human observers were still able to hold color constancy only from a simple pattern of colors.

In addition to the simplification of surface properties, there are at least two limitations regarding illuminant properties in this study. First, our test illuminants always changed along the blue-yellow direction. It would be interesting to test whether human color constancy still holds for non-black-body locus illuminants (e.g. cyan or magenta illuminant) to which we are not exposed in daily life. Interestingly Delahunt and Brainard (2004) showed that our color constancy is not impaired under an atypical illuminant, suggesting our internal assumption about illuminant color is not constrained along the blue-yellow axis. Second, all experimental scenes were uniformly illuminated by a single illumination. In contrast, objects placed in the real world tend to receive incident lights from every direction and the spectral distribution of the lights may change from one direction to another. For example, in sunny outdoor scenes light from above tends to be sunlight or skylight but the object also receives light from below dominated by a secondary reflection from other objects in the scene. Recent studies indeed have shown that natural scenes have a significant amount of directional spectral variation (Morimoto et al., 2019). For a scene illuminated by a single light source, as shown in Figure 1, a single optimal color distribution shows a completely representation of the surface color gamut. However, for a scene under multiple illuminants the gamut accordingly expands, and thus our model also needs to consider more than one optimal color distribution. In general estimation of multiple illuminations increases the complexity of the color constancy problem and thereby inflates computational cost, and therefore our model may also suffer because there are too many combinations of optimal color distributions to fit. In recent years an increasing body of research has investigated the influence of directional-dependent illumination on human color constancy or other functions of color vision (Fleming, Dror and Adelson, 2003; Doerschner, Boyaci, and Maloney, 2004; Morimoto and Smithson, 2018). Our present study suggested that the optimal-color-based explanation reasonably works when there is only one illuminant in a scene. Testing the limitation of human color constancy under multiple illuminants allows us to see the degree to which our proposed model can be applied.

Our model has some parallels with the idea that visual system uses statistical regularities hidden in natural scenes which can act as internal references to calibrate our perception (Gilchrist et al, 1999; Lee and Smithson, 2012). For example, our white point settings when no context is presented seems to spread along the chromatic variation of natural scenes, implying mechanisms to normalize color appearance to scene statistics (Bosten, Beer and MacLeod, 2015). There are also a number of empirical data that suggest the influence of natural scene statistics on color vision mechanisms such as chromatic adaptation, color discrimination, and color constancy (Webster and Mollon, 1997; MacLeod, 2003; Webster 2011). Moreover, there is an evidence that our unique hue points can be recalibrated following the exposure to the manipulated chromatic signals (Neitz et al., 2002) or the seasonal change of chromatic statistics (Welbourne, 2015). DeLawyer et al. (2018) showed that the amount of S-cone excitation could be a cue to judge whether a color change stems from an illuminant change or a surface change. Major findings in the present study is that we can account for human observers’ estimation of illumination if we assume that the visual system has full access to the distribution of optimal colors. However, how much are these assumptions valid? How plausible is it for us to know physical limit of colors? Rather it seems to be an open question still as to whether the human observers are able to access the shape of the optimal color distribution through the observation of statistical regularities in natural scenes. Figure 1 shows that when we plot 49,667 natural objects at once, it very much resembles the shape of the optimal color distribution. This indicates that if we integrate the luminance vs. chromaticity distribution over a long period, it may be possible for us to learn where the physical limits are. We believe that the observer’s internal assumption about the physical limit of surface color can be visualized by measuring luminosity threshold over various chromaticities. Our unpublished data (Uchikawa et al., 2001; Uchikawa, Fukuda, and Morimoto, 2017) indicated that the locus of luminosity thresholds resembles the luminance of optimal colors. Additionally, Speigle and Brainard (1996) suggested that it is possible to relate luminosity thresholds to the upper luminance boundary estimated from natural objects’ luminance distribution, which agrees with the optimal color distribution as shown in Figure 1. Recent studies also have begun to show success of such learning-based approach in other research domains (Fleming and Storrs, 2019). For complex functions such as color constancy where complete implementation requires solving inverse optics, a learning-based strategy provides a powerful alternative. Thus, it may be plausible for our visual system to learn and utilize the optimal color distribution to implement a mathematically challenging constancy mechanism. The extension of research in this direction might yield exciting insights to long-standing questions in the field of color vision.

## Acknowledgement

This work was supported by JSPS KAKENHI Grant Number JP19K22881, JP17K04503 and 26780413. TM is supported by a Sir Henry Wellcome Postdoctoral Fellowship awarded from the Wellcome Trust. The authors are grateful to Tanner DeLawyer for the correction of grammatical errors in main text.

